# Tumor Protein D54 (TPD54) regulates intracellular protein trafficking, cellular function and disease progression in melanoma

**DOI:** 10.64898/2026.05.07.721771

**Authors:** Michael Ortiz, Charlie B. Ffrench, Stuart Webb, John Toubia, Nathalie Nataren, Emma L. Dorward, Kay K. Myo Min, Ana Lonic, Peer Arts, Michaelia P. Cockshell, My G. Mahoney, Michael S. Samuel, Lisa M. Ebert, Yeesim Khew-Goodall, Claudine S. Bonder

## Abstract

To facilitate survival, migration and evasion of immune surveillance, cancer cells tightly coordinate the synthesis and trafficking of a diverse repertoire of proteins to their cell surface and the surrounding tumor microenvironment. A key mechanism underlying this process is the intracellular membrane trafficking pathways, including vesicular transport systems. There remains a paucity of mechanistic insight into the regulatory components that mediate nascent protein trafficking and their dysregulation in cancer. Herein, we investigate Tumor Protein D54 (TPD54) as a central regulator of intracellular protein transport that is exploited by melanoma cells to promote disease progression. Integrative analyses of patient-derived tumor tissue specimens show that the expression of *TPD52L2* (the gene encoding TPD54) is frequently overexpressed in melanoma and correlates with adverse clinical outcomes, including reduced responses to immune checkpoint blockade. Mechanistic investigations further revealed that TPD54 maintains Golgi integrity and orchestrates trafficking of early endosomes, anterograde vesicles and extracellular vesicles. Functionally, TPD54 augments the secretion of pro-cancerous cytokines, increases the cell surface expression of adhesion-signaling receptors (e.g. integrin-β1 and desmoglein-2), promotes melanoma cell migration and elevates their capability to undergo vasculogenic mimicry. Targeting *TPD52L2* in two mouse models of melanoma (B16-F10 and HCmel12) showed significant attenuation of tumor growth, disrupted tumor vasculature, enhanced anti-tumor immunity with infiltration of CD8^+^ T cells and reduced metastatic disease. Collectively, these findings establish TPD54 as a critical and previously underappreciated regulator of protein trafficking in cancer cells that directly contributes to disease progression and highlights its potential as a novel therapeutic target to combat melanoma.

## Introduction

Melanoma remains the most lethal form of skin cancer, accounting for approximately 80% of global skin cancer-related deaths (1, 2). Although therapeutic advances, including BRAF/MEK inhibitors and immune checkpoint blockade (e.g., anti-CTLA-4 and anti-PD-1) have been transformative for a subset of patients, inherent or acquired resistance remains a significant problem (3–6) and for these patients the overall 5-year survival rate for metastatic disease is approximately 30% (1, 7). Together with the high metastatic propensity of melanoma, these limitations underscore the continued need for novel therapeutic strategies, guided by a more comprehensive understanding of the molecular and cellular mechanisms driving melanoma progression.

A major contributor to melanoma pathogenesis is the tumor microenvironment (TME), a dynamic and heterogeneous ecosystem comprised of cancer cells, stromal cells, immune cells, and acellular components of the extracellular matrix (8, 9). The TME actively fosters cancer progression via immunosuppressive signaling events, recruitment of pro-tumorigenic immune cells and exclusion or functional impairment of anti-tumor immune cells (10). In particular, cytotoxic CD8⁺ T cells and natural killer (NK) cells frequently exhibit limited infiltration and persistence within the tumor mass. This restricted access is due, at least in part, to physical and molecular barriers, including abnormal tumor vasculature (10–14). To establish and maintain these immunoregulatory mechanisms, cancer cells dynamically reprogram intracellular vesicle trafficking and protein transport pathways (8, 15–18).

An emerging regulator of intracellular protein trafficking is Tumor Protein D54 (TPD54). As one of four members of the tumor protein D52-like family (TPD52-TPD55), TPD54 upholds the membrane integrity of the Golgi apparatus and engages in intracellular vesicle trafficking pathways; particularly anterograde transport (the forward movement of proteins/molecules from the inner cell body to the plasma membrane), as well as vesicle recycling (19–21). In HeLa cells, TPD54 forms complexes with Rab GTPases and associates with intracellular nanovesicles where it regulates the trafficking of cargo proteins, including integrin α5β1 (20). Notably, *TPD54* gene expression is elevated in tumor tissues from the breast, brain, lung, and kidney; and is linked to key malignant behaviors including cancer cell proliferation, migration and invasion (19, 20, 22–33). A recent gene set enrichment analysis (GSEA) of patients with lung adenocarcinoma showed that *TPD54* was associated with cell cycle and immune-regulation-related pathways, such as “G1/S Transition”, “Cell Cycle”, “Adaptive Immune System” and “Innate Immune System” (34). Despite these observations, there remains a paucity of understanding on the molecular mechanisms underlying TPD54-mediated cancer progression.

Herein we directly compare the expression of *TPD52L2* and its family members in healthy skin and melanoma and align with a survival profile. We examine partnerships between TPD54 and intracellular trafficking proteins (Rab GTPases), identify proteins reliant on TPD54 for cell surface expression and secretions, and assess how this small protein (∼20 kDa) influences melanoma cell function and disease progression *in vivo*. We investigate the influence of TPD54 on anti-tumor immunity and compare TPD54 function between melanoma and healthy cells. Collectively, these insights substantially advance our current knowledge of TPD54-mediated vesicular trafficking and establish its potential relevance for melanoma prognostication and therapeutic intervention.

## RESULTS

### *TPD52L2* expression in melanoma tissues is associated with high-risk disease

To investigate the role of TPD54 in melanoma, we analyzed transcriptomic data from the NCBI Gene Expression Omnibus (GEO) comprising primary malignant melanoma tumors (n=46), benign nevi (n=18), and normal skin tissues (n=7). Microarray profiling revealed *TPD52L2* expression to be significantly higher in malignant melanoma compared to both normal skin and benign nevi **(Figure 1a)**. The elevated expression of *TPD52L2* in melanoma tissues differed from other family members, with loss of *TPD52L1* (which encodes TPD53) associated with malignant melanoma **(Figure 1b)** and no associations identified with *TPD52* expression levels **(Figure 1c)**, raising the possibility that *TPD52L2* may disproportionately influence the molecular pathways governing melanoma progression. As TPD55 (encoded by *TPD52L3*) is uniquely restricted to the testis (19) it was not included in the study.

**Figure 1:**
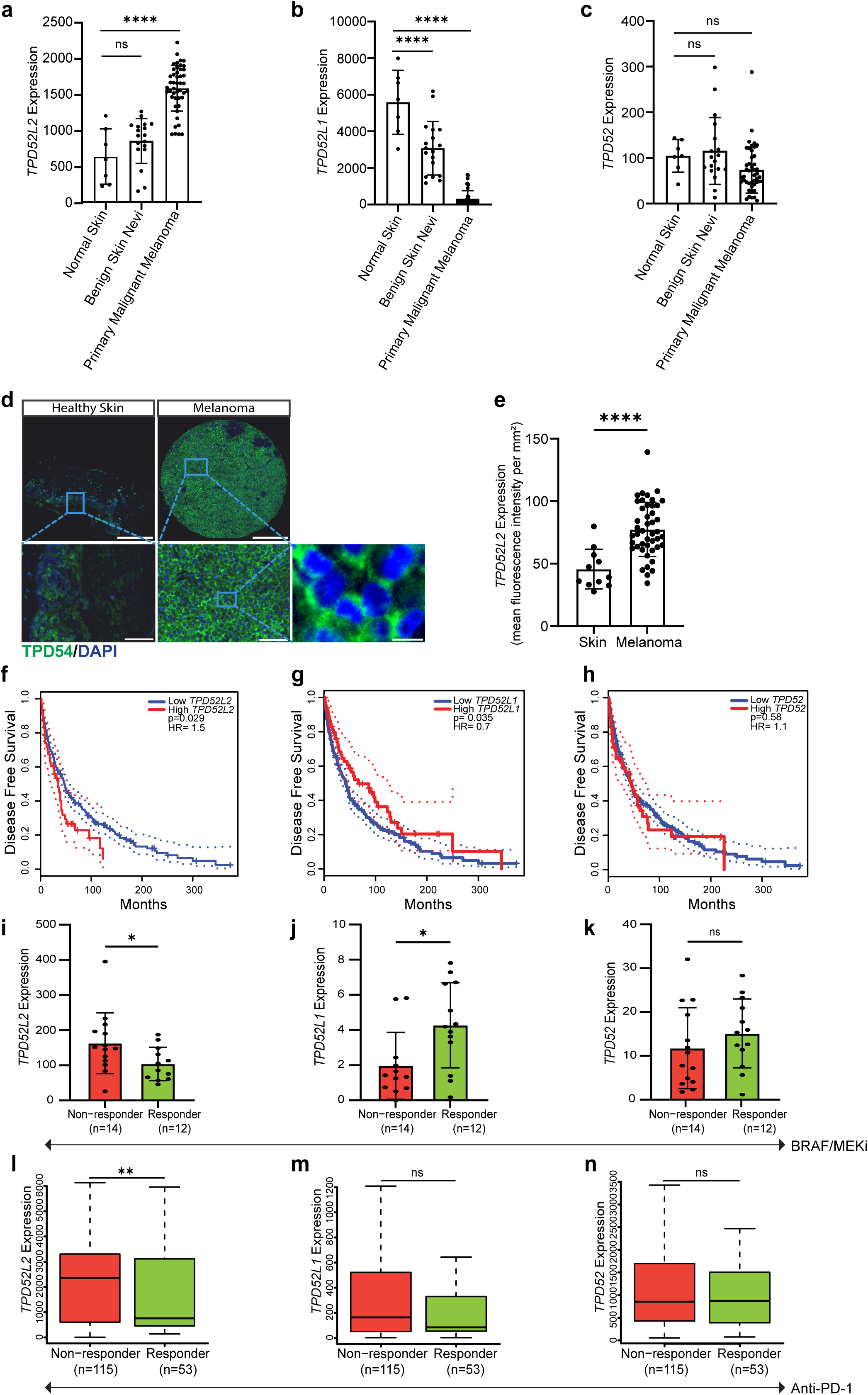
Expression of *TPD52L2*, *TPD52L1*, *TPD52* in human healthy skin and melanoma tissue, survival outcomes and treatment response. Comparative analysis of **a)** *TPD52L2*, **b)** *TPD52L1* and **c)** *TPD52* expression in normal skin (n=7), benign skin nevi (n=18) and primary malignant melanoma (n=46) within GEO dataset (GEO accession number: GSE3189). **d)** Representative image of IF staining for TPD54 on TMAs containing healthy skin (n=11) and melanoma (n=45), scale bar=100 µm. **e)** From **d** a quantification of the number of TPD54+ cells per core. Data is presented as mean ± SEM. Kaplan–Meier plots indicating the DFS of melanoma patients with low (blue n=413) and high (red n=413) f) *TPD52L2*, g) *TPD52L1* and h) *TPD52* levels from TCGA database. Comparative gene expression of i) *TPD52L2*, j) *TPD52L1* and k) *TPD52* in melanoma responder vs non-responders to BRAF + MEK inhibitor (responder n=12-13, non-responders n=12-14) from the GEO dataset (GEO accession number: GSE196434). Comparative gene expression of l) *TPD52L2*, m) *TPD52L1* and n) *TPD52* for PD-1 inhibitor (Nivolumab) resistance (non-responder n=115, responder n=53) in melanoma patients. Statistical analysis using one-way ANOVA method **(a-c)**, log-rank test **(f-h)** or unpaired *t-*test **(d,e,i-n)**, * = p<0.05, **** = p<0.0001.

Next, TPD54 protein expression was assessed by immunofluorescence (IF) analysis of a tissue microarray (TMA) containing cores from healthy skin tissue (n=11 donors) and melanoma (n=45 donors). This analysis revealed showing higher TPD54 protein levels in the tumor tissue compared to normal skin **(Figure 1d/e)**.

To determine whether *TPD52L2* levels correlate with risk, patient tumors within the GEPIA melanoma dataset (35) were stratified via median split into *TPD52L2* high- and low-expression groups (n=413 per group). High *TPD52L2* expression correlated with significantly worse disease-free survival (DFS) for patients when compared to those with tumors of low *TPD52L2* expression (p = 0.029; HR = 1.5; **Figure 1f**). Consistent with the gene expression profiling above, low *TPD52L1* expression correlated with worse DFS (p = 0.035; HR = 0.7; **Figure 1g**) while *TPD52* levels did not associate with DFS (p = 0.58; HR = 1.1; **Figure 1h**).

Building on the potential of *TPD52L2* as a prognostic biomarker in melanoma, emerging evidence suggests that abnormal gene expression profiles may also serve as predictors of therapeutic response. First, we evaluated *TPD52L2* expression levels in melanoma tumor samples from patients treated with inhibitors to BRAF + MEK (dataset GSE196434, (36)) classifying patients as responders (partial response or complete response with progression-free survival (PFS) >12 months) or non-responders (<12 months of PFS). *TPD52L2* expression was significantly higher in the tumors of patients of BRAF + MEKi non-responders compared to responders (**Figure 1i**). In contrast, more durable responses to BRAF + MEK inhibition were observed in patients whose *TPD52L1* levels were high (**Figure 1j**), while *TPD52* expression was not informative of responsiveness to this therapy regime (**Figure 1k**). Next, using data from the ROCplot platform (37) we investigated *TPD52L2* expression in tumor tissues from melanoma patients (primary and metastatic, pre-treatment) who went on to be treated with anti-PD-1 (nivolumab). **Figure 1l** shows that *TPD52L2* expression was significantly higher in non-responders compared to responders, while the levels of neither *TPD52L1* or *TPD52* were informative of anti-PD-1 responsiveness (**Figures 1m,n**). Consistent with these findings, analysis of overall survival (OS) derived from the TCGA, GEO and EGA databases, revealed that elevated *TPD52L2* expression (in pretreatment samples) was associated with poorer outcomes for melanoma patients administered with either anti-PD-1 (nivolumab or pembrolizumab) or anti-CTLA4 (ipilimumab) (**Supplementary Figure S1a, b**). Interestingly, for patients treated with anti-PD1, high *TPD52L1* expression also correlated with reduced OS (**Supplementary Figure S1a**). *TPD52* expression was again non-informative for OS with this therapy (**Supplementary Figure S1a, b**). Taken together, this transcriptomics data suggests that high expression of *TPD52L2* in melanoma is associated with high-risk disease, including treatment resistance.

### TPD54 mediates vesicle trafficking and secretion from melanoma cells

To investigate the role of TPD54 in melanoma, we first examined its subcellular distribution with initial focus on the Golgi apparatus (19, 20). Immunofluorescence analysis and confocal microscopy of human melanoma cell lines (C32, CHL-1, and HT-144) revealed low diffuse cytoplasmic distribution of TPD54 throughout the cells with notable co-localization to the GM130+ stained Golgi matrix **(Figure 2a, b)**. To examine whether TPD54 is important for Golgi integrity, siRNA-mediated knockdown of *TPD52L2* was undertaken **(Figure 2c, d)** and assessed for trans-Golgi network (TGN) morphology by TGN46 staining. Compared to control cells (siCTRL), TPD54-depleted melanoma cells (si*TPD52L2*) exhibited a fragmented and swollen TGN morphology **(Figure 2e)**, with a significantly higher Golgi area **(Figure 2f)**, indicative of structural disassembly (38–42). To investigate whether this disruption of Golgi structure affected intracellular trafficking events, members of the Rab GTPase family (key regulators of distinct stages of vesicle transport (43–46)) were employed as markers of intracellular vesicles. *TPD52L2*-knockdown in the three melanoma cell lines did not alter distribution of markers for early endocytosis (Rab5) or late endosome/lysosomes (Rab7) **(Figure 2g, h)**. In contrast, we observed a greater dispersal and expression of markers of early endosomes (Rab4), cargo transport from the Golgi apparatus to early endosome (Rab9) and anterograde vesicle trafficking to the cell membrane (Rab10) **(Figure 2i-k)**. Taken together, these data suggest that depletion of TPD54 in melanoma causes trafficking delays or vesicle stalling, which is consistent with impaired Golgi function (17, 19, 38–42).

**Figure 2:**
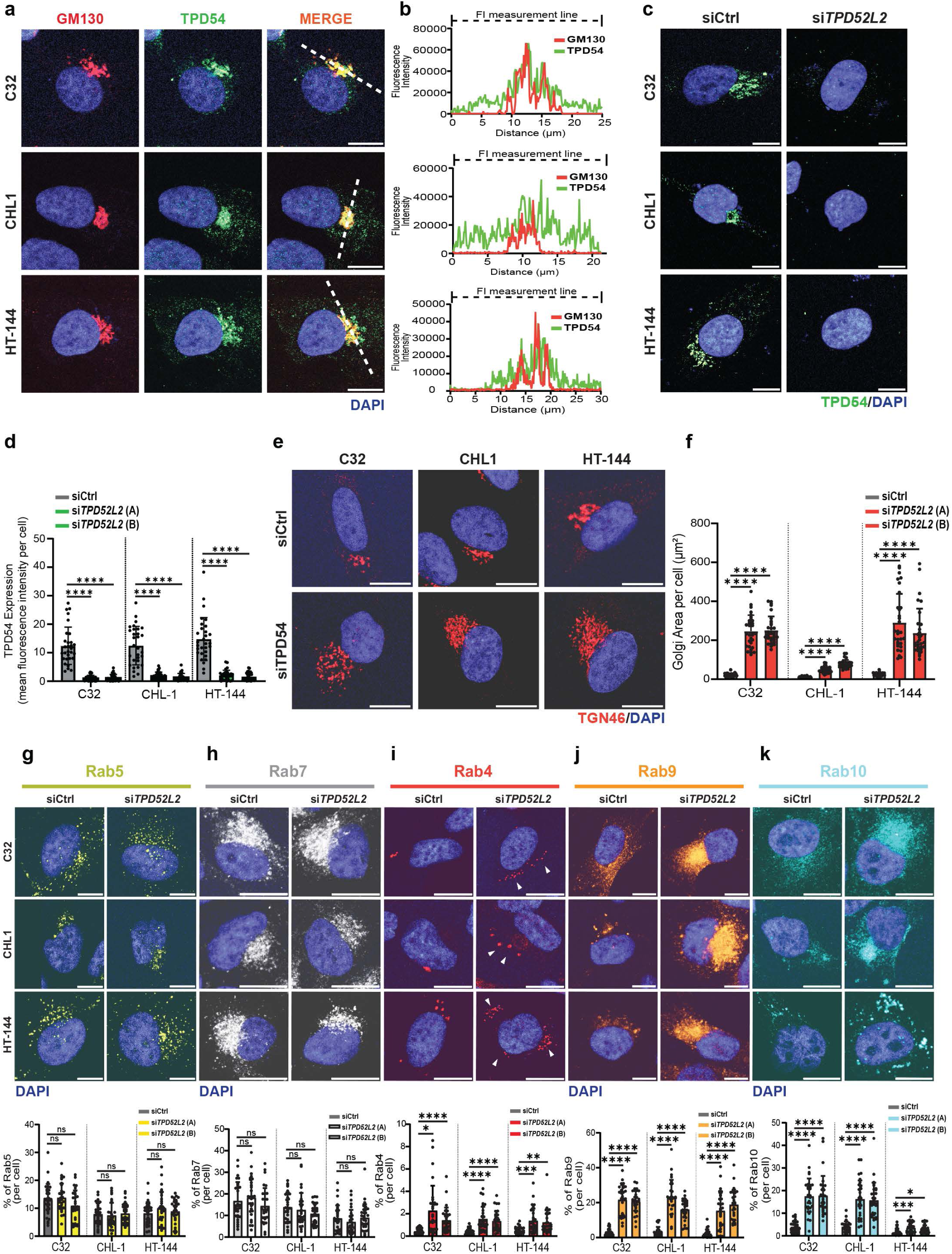
TPD54 expression in intracellular vesicles of melanoma cells. **a)** IF images and **b)** distribution plots of TPD54 (green) in melanoma cells (C32, CHL1, HT-144) relative to and Golgi positioning (GM130, red)). Scale bar = 10 μm. The fluorescent intensity profile was measured on the merged image (dashed line). **c)** IF images of TPD54 (green) expression in response to knockdown of *TPD52L2* in melanoma cells, nucleus (blue). Scale bar, 10 μm **d)** Quantification of TPD54 expression in melanoma cells with *TPD52L2* KD (si*TPD52L2* (A) or si*TPD52L2* (B)) via fluorescent intensity (n=3 individual experiments/30 cells per group). **e)** IF images of trans-Golgi network (TGN46, red) in response to the downregulation of *TPD52L2* in melanoma cells, nucleus (blue). Scale bar=10 μm. **f)** Quantification of TGN46+ Golgi area/per cell (μm^2^) (n=3 individual experiments/30 cells per group). (**g-k**) IF images and quantification of % Rab+ proteins in response to *TPD52L2* KD in melanoma cells. **g)** Rab5 (yellow), **h)** Rab7 (white), **i)** Rab4 (red), **j)** Rab9 (orange), **k)** Rab10 (cyan), nucleus (blue). Scale bar=10 μm, (n=3 individual experiments/30 cells per group). Data is expressed as mean±SEM, ns= >0.05, * = p<0.05, ** = p<0.01, *** = p<0.001 and **** = p<0.0001 using one-way ANOVA analysis.

As Rab9 and Rab10 have established roles in extracellular vesicle (EV) trafficking and secretion (47), we investigated whether TPD54 might regulate EV biogenesis. EVs were isolated from the culture media of three melanoma cell lines without and with *TPD52L2*-knockdown, lysed and examined for purity via immunoblotting for CD81 (EV membrane maker) and ALIX (EV cytosolic maker). In addition, GM130, a non-EV marker was included as a negative control to ensure the specificity of EV isolation and to rule out contamination by non-vesicular cellular structures **(Figure 3a)**. EV lysates of all melanoma cell lines (C32, CHL-1 and HT-144) with *TPD52L2* knockdown had significantly lower levels of ALIX (EV cytosolic marker) and CD81 (EV membrane marker) **(Figure 3a, b)** compared to controls. These findings were corroborated by nanoparticle tracking analysis (NTA), which showed significantly fewer EV particles in the 50–200 nm diameter range, consistent with the size of exosomes, in TPD54-depleted samples relative to controls **(Figures 3c, d and Supplementary video S1, S2)**.

**Figure 3:**
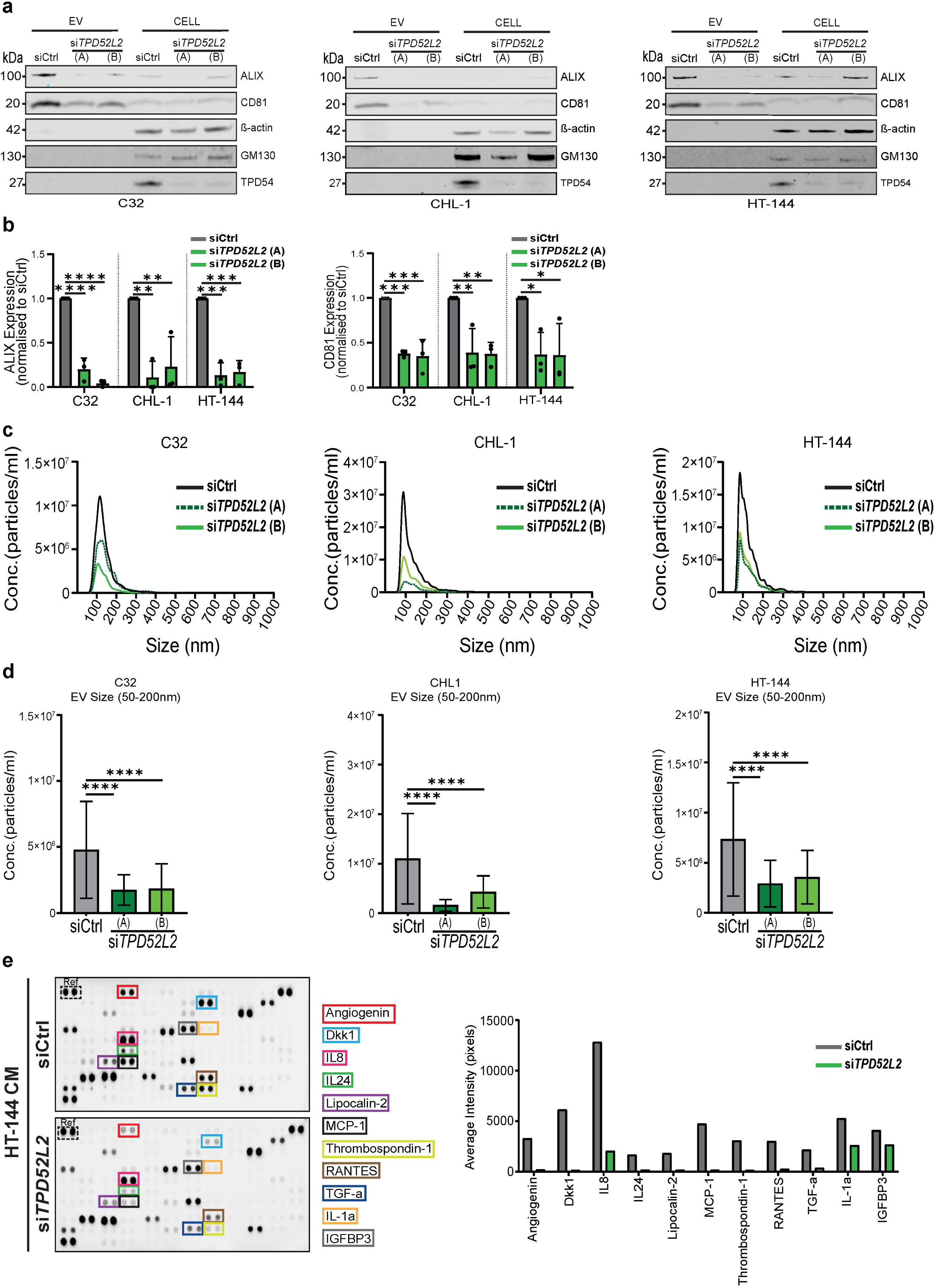
TPD54 and extracellular vesicle secretion by melanoma cells. **a)** Immunoblot of extracellular vesicles (EVs) and cell lysates of melanoma cells (C32, CHL1, HT-144) with *TPD52L2* KD **b)** Protein quantification of ALIX and CD81 (EV markers) (n=3 individual experiments). **c)** Nanoparticle-tracking analysis (NTA) of EVs in response to *TPD52L2* downregulation. Histogram showing the EV particle-size distribution (size in nanometers) (n=3 individual experiments). **d)** EV size change in response to TPD54 down regulation (50nm-200nm size range). **e)** Representative images and analysis of proteome profiling array on conditioned media (CM) from HT-144 cells without or with *TPD52L2* knockdown. Data are presented as mean ± SEM. Analysis via one-way ANOVA method (**b-i, k**) to compare against siCtrl controls. * = p<0.05, ** = p<0.01, *** = p<0.001 and **** = p<0.0001.

To further characterize the impact of TPD54 on broader secretory pathways in melanoma, a cytokine profiling array was conducted on media conditioned by HT-144 melanoma cells with and without *TPD52L2* knockdown. The levels of several secreted factors, including angiogenic regulators (angiogenin, dickkopf-related protein 1 (DKK1)), immune modulators (MCP-1, IL-8, IL-24), and inflammatory cytokines and growth factors (TNF-α, thrombospondin-1) were diminished in media conditioned by HT-144 melanoma cells in which *TPD52L2* had been knocked down, compared to media conditioned by control HT-144 melanoma cells **(Figure 3e)**. Taken together, these data suggest that TPD54 mediates major biological pathways that influence the local tumor microenvironment.

### TPD54 facilitates the transport to the cell surface of proteins known to mediate tumor-promoting signaling pathways

To investigate whether TPD54 also regulates the mobilization of proteins to the plasma membrane, we next examined the cell surface localization of two proteins documented to promote melanoma, a cadherin (desmoglein-2 (DSG2)(48, 49)) and an integrin (i.e. integrin-β1, (50–56)). We first assessed the interaction between TPD54 and DSG2 via confocal microscopy and observed broad expression of DSG2 throughout the cell as well as defined areas of co-localization with TPD54 in the three melanoma cells lines **(Figure 4a, b)**. Co-immunoprecipitation with an antibody against TPD54 followed by immunoblotting for both TPD54 and DSG2 confirmed that TPD54 and DSG2 co-occurred in a protein complex **(Figure 4c)**. Furthermore, TPD54 appears to have a role in trafficking DSG2 in melanoma cells, as cells in which *TPD52L2* has been knocked down exhibited accumulation of DSG2 within cytoplasmic compartments, across all three cell lines **(Figure 4d)**. Consistent with this observation, immunoblotting of whole cell lysates revealed higher total DSG2 protein levels in TPD54-depleted cells **(Figure 4e)**. To directly assess the possible impairment of DSG2 trafficking to the cell surface, we utilized an acid-wash recovery assay wherein flow cytometric analysis determined that removal of DSG2 from the cell surface is replenished within six hours **(Supplementary Figure S2a)**. Analysis of the melanoma cell lines with *TPD52L2*-knockdown at six hours (T=6) post acid-stripping revealed a significant inhibition of DSG2 resurfacing **(Figure 4f)**. Consistent with our observations for DSG2, TPD54 also co-localized with a proportion of integrin-β1 within melanoma cells **(Figure 4g, h)** and *TPD52L2* knockdown resulted in an intracellular accumulation of integrin-β1; further supporting impaired expression of DSG2 and integrin-β1 on the cell surface **(Figure 4i)**.

**Figure 4:**
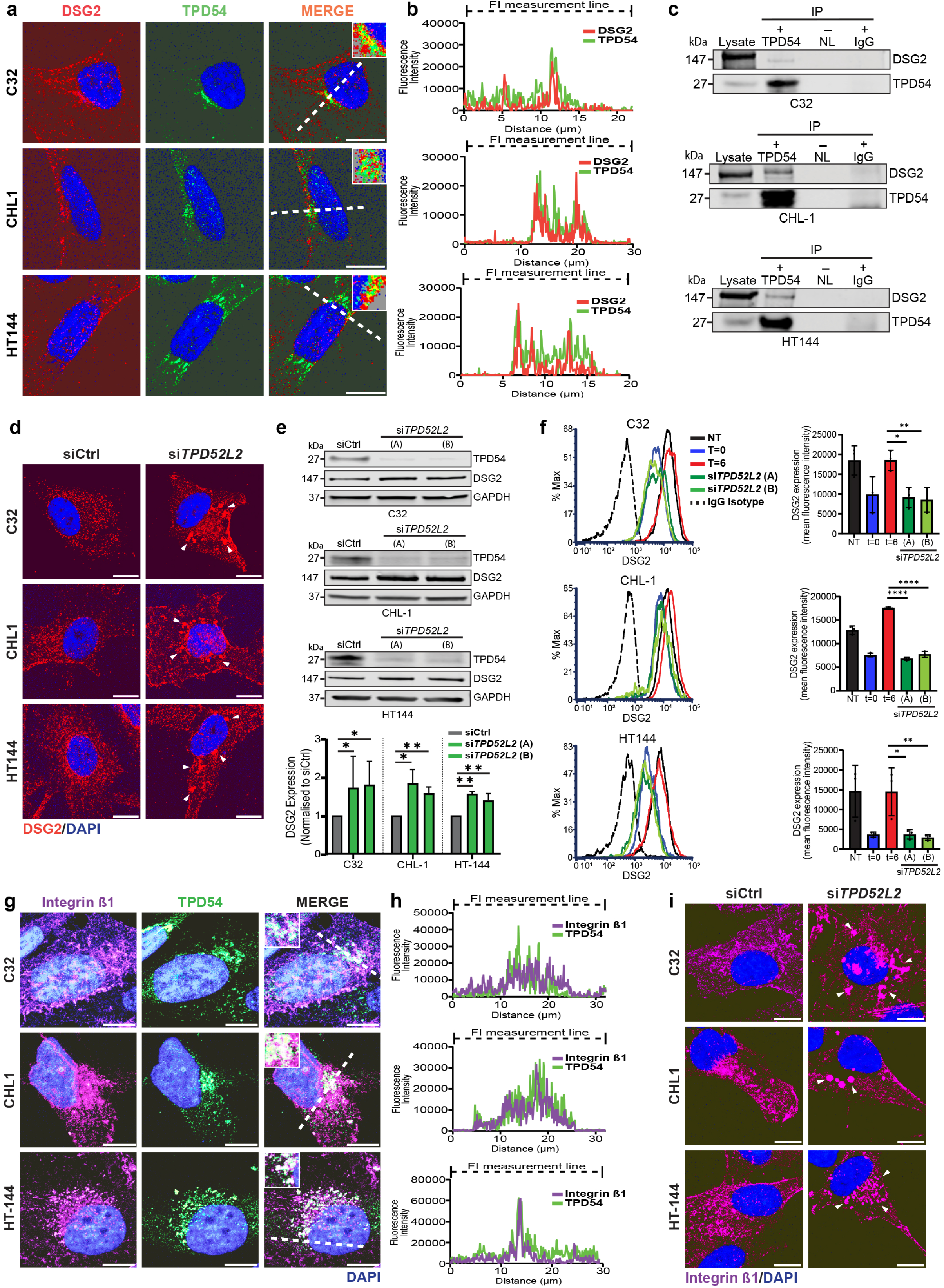
TPD54 mediates protein trafficking and expression of DSG2 and integrin β1 in melanoma. **a)** IF images of TPD54 (green) and DSG2 (red) in melanoma cells (C32, CHL1, HT-144) Scale bar = 10 μm. **b)** distribution plots. The fluorescence intensity profile representing the dashed line in the previous image=. **c)** Immunoprecipitation of TPD54 from melanoma cells followed by immunoblot for TPD54 and DSG2. **d)** Representative IF images of DSG2 localization in melanoma cells without or with *TPD52L2* KD. Scale bar = 10 μm. **e)** Immunoblot of DSG2 in melanoma cells ± *TPD52L2* KD (n=3 per cell line). **f)** Acid stripping and resurfacing of DSG2 in melanoma cells ± *TPD52L2* KD with flow cytometry histograms (left) and quantified data from n=3 independent experiments. **g)** Representative IF images of integrin-β1 (purple) and TPD54 (green) in the melanoma cells. Scale bar = 10 μm. **h)** distribution plots representing the dashed line in the previous image. **i)** Representative IF images of integrin-β1 (purple) localization in melanoma cells ± *TPD52L2* KD with nuclei DAPI stained (blue). Scale bar = 10 μm. Data are expressed as mean±SEM, ns= >0.05, * = p<0.05, ** = p<0.01, *** = p<0.001 and **** = p<0.0001 using one-way ANOVA method.

### TPD54 promotes melanoma cell migration and vasculogenic mimicry

We next sought to determine whether TPD54 influences melanoma cell functions, such as cell survival and migration. To address this, C32, CHL-1, and HT-144 melanoma cells without and with *TPD52L2* knockdown were examined at 72 hours for apoptotic (Annexin-V+) and necrotic (7AAD+) markers via flow cytometry. TPD54-depletion did not affect cancer cell survival **(Figure 5a)**. Given the disruption in integrin-β1 and Rab4 trafficking observed following *TPD52L2* knockdown (both of which are critical mediators of cell migration (20, 57–59)) we next assessed whether TPD54 modulates melanoma cell migration. Transwell migration assays showed substantially lower migratory capacity across all TPD54-depleted melanoma cell lines compared to controls over a 24–48-hour period **(Figure 5b, c)**. Aggressive melanoma cells undergo vasculogenic mimicry (VM) to form their own vessel-like structures devoid of endothelial cells to promote cancer progression (49, 60–62). As both DSG2 and integrin-β1 regulate VM in melanoma (49, 63), we examined whether TPD54 might promote VM. Indeed, **Figure 5d-e** shows that TPD54-deficient melanoma cells fail to form VM networks *in vitro*, instead aggregating into disorganized clusters **(Sup Videos 3-8)**.

**Figure 5:**
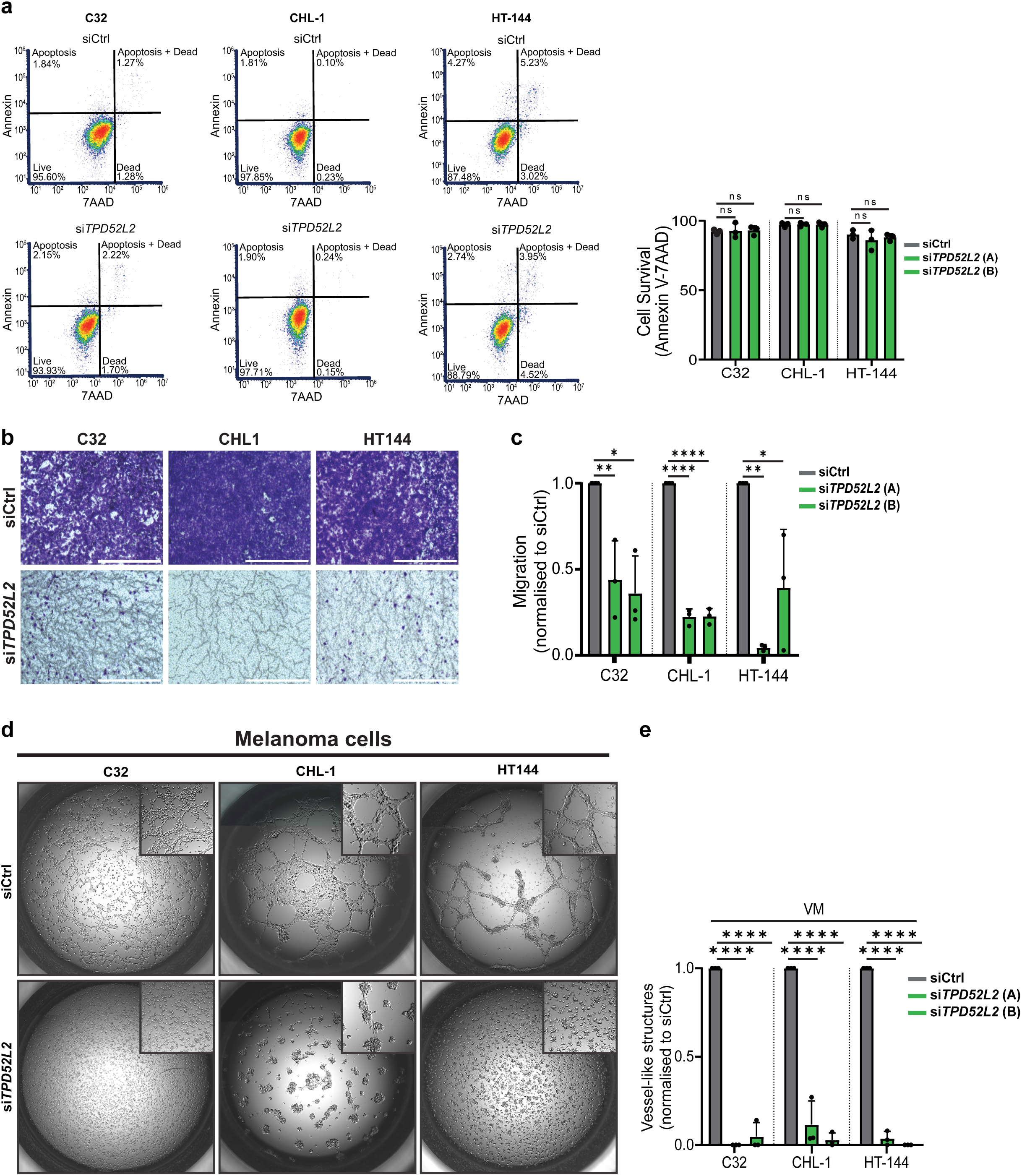
TPD54 supports melanoma cell migration and vasculogenic mimicry. **a)** Cell survival analysis via staining for Annexin V (apoptosis) and with 7-AAD (cell death), flow cytometric analysis of melanoma cells (C32, CHL1, HT-144) ± *TPD52L2* KD. Data represent mean ± SEM and analyzed via one-way ANOVA. **b)** Melanoma cell migration following *TPD52L2* KD was assessed via a Transwell migration assay towards 10% FBS. Representative images of migrated cells stained with crystal violet. **c)** Quantification of migrated cells from (b) using ImageJ analysis. Data represent n = 3 independent experiments. **d)** Representative images of vasculogenic mimicry by melanoma cell lines (C32, CHL-1, HT-144) ± *TPD52L2* KD and between 12-24hr after seeding cells onto Matrigel. **e)** Quantification of VM structures from **d),** per cell line. Data are presented as mean ± SEM and analyzed via one-way ANOVA (d, f, h) compare siCtrl controls. ns= >0.05, * = p<0.05, ** = p<0.01 and **** = p<0.0001

### Cellular functions of TPD54 vary between healthy and cancerous cells

We next examined whether TPD54 had functions in healthy cells that were similar to those demonstrated above. First, we applied the experimental pipeline detailed above on a transformed human bone marrow derived endothelial cell line (TrHBMEC, hereafter labelled as BMEC) (64, 65). Immunofluorescence analysis was used to first demonstrate successful depletion of TPD54 in BMEC **(Figure 6a)**. As in melanoma cells, TPD54 also co-localizes with the Golgi apparatus in endothelial cells **(Figure 6b).** But in contrast to melanoma cells, *TPD52L2* depletion in BMEC did not affect TGN morphology **(Figure 6b-d)**. Also, unlike melanoma cells TPD54 did not co-immunoprecipitate with DSG2 in BMEC suggesting that they did not occur in complex with each other **(Figure 6e)** and accordingly, *TPD52L2* knockdown did not impair DSG2 intracellular trafficking **(Figure 6f-h)**. Furthermore, *TPD52L2* knockdown in BMEC did not impair *in vitro* angiogenesis **(Figure 6i, j & Sup Videos 9-10)**. Similar observations were made in HaCaT keratinocytes, where *TPD52L2* knockdown did not cause fragmentation of the Golgi (**Figure 6k, l**).

**Figure 6:**
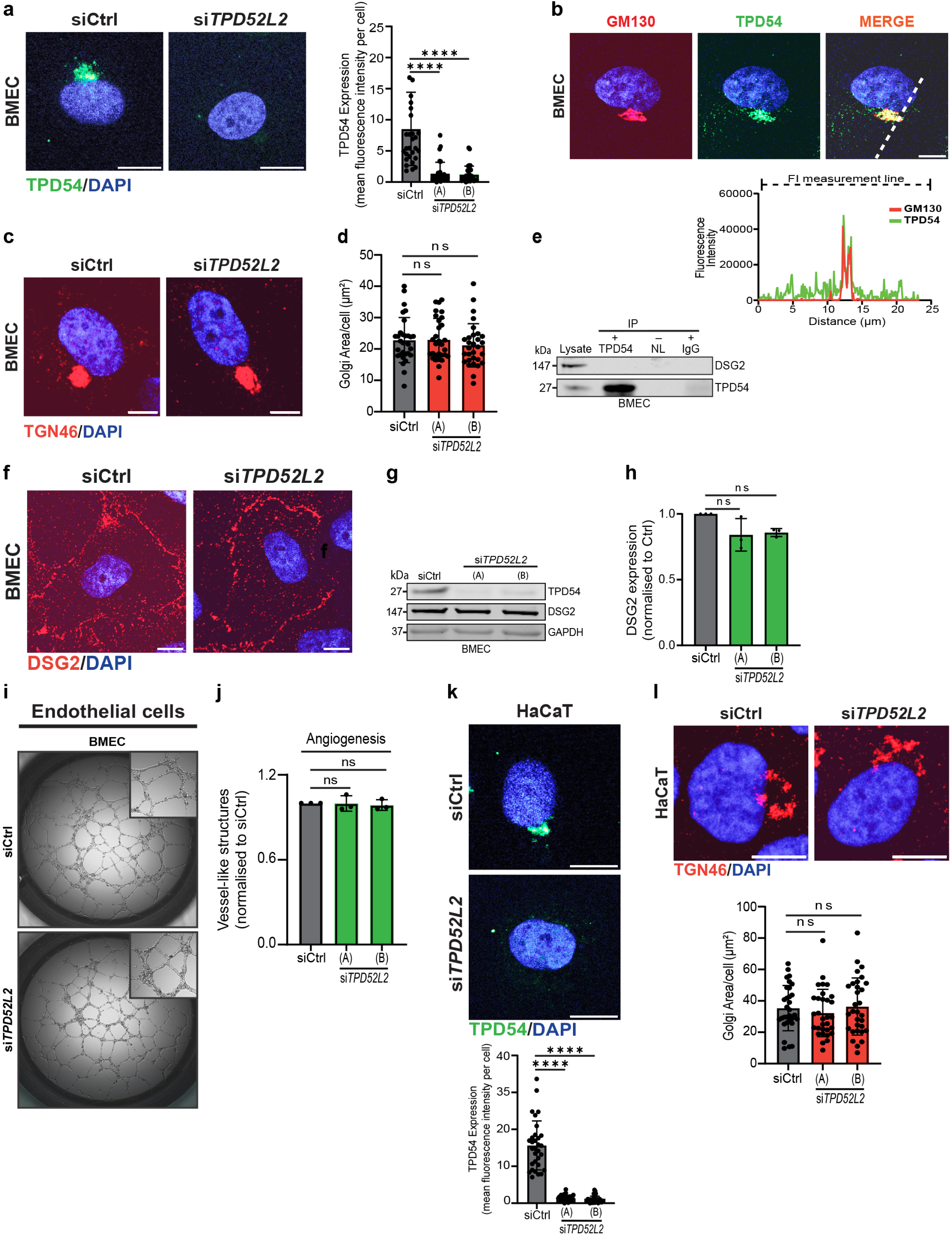
TPD54 expression and function in endothelial cells and keratinocytes. **a)** Representative IF image of TPD54 (green) and nuclei (DAPI, blue) in BMEC and quantification of *TPD52L2* KD (n=3 individual experiments/30 cells per group). Scale bar = 10 μm. **b)** Representative IF images of TPD54 (green), Golgi (GM130, red) and nuclei (DAPI, blue) with distribution profile (dotted line). Scale bar = 10 μm. **c)** Representative IF images of trans-Golgi network (TGN46, red) and nucleus (DAPI, blue) in BMEC ± *TPD52L2* KD and **d)** quantification of Golgi area/per cell (μm^2^) (n=3 individual experiments/30 cells per group). **e)** Immunoprecipitation of TPD54 from BMEC, with immunoblot for TPD54 and DSG2, representative of three independent experiments. **f)** Representative IF images of DSG2 (red) and nuclei (DAPI, blue) expression in BMEC ± *TPD52L2* KD. Scale bar = 10 μm. **g-h)** Representative immunoblot of DSG2 in BMEC ± *TPD52L2* KD, GAPDH as loading control (n=3 independent experiments). **i)** Representative photomicrographs and **j)** quantification of angiogenesis by BMEC ± *TPD52L2* KD 24hr after seeding on Matrigel. **k)** Representative IF images of TPD54 (green) and nuclei (DAPI, blue) ± *TPD54* KD in HaCaT keratinocyte cells, with quantification of TPD54 expression via fluorescence intensity (n=3 individual experiments/30 cells per group). Scale bar=10 μm. **l)** Representative IF images of trans-Golgi network (TGN46, red) in HaCaT keratinocytes ± *TPD52L2* KD, nucleus (DAPI, blue). Scale bar=10 μm. Quantification of Golgi area/per cells (μm^2^) (n=3 individual experiments/30 cells per group). Data are presented as mean ± SEM and analyzed using one-way ANOVA to compare against siCtrl. ns= >0.05 and **** = p<0.0001

To address whether the disparities in TPD54 function between cancerous cells and healthy cells could be attributable to differing expression levels, we compared protein quantities of TPD52, TPD53 and TPD54 across all the cell lines. **Figure 7a** shows that all TPD family members are more abundant in melanoma cells (C32, CHL-1, HT-144) compared to BMEC endothelial cells. In HaCaT keratinocytes, protein levels of TPD52 and TPD53 were most abundant when compared to melanoma **(Figure 7b)**. To investigate the potential for compensation across TPD family members, *TPD52L2* was depleted in the cancerous and healthy cells and caused a significant increase in both TPD52 and TPD53 protein levels in melanoma **(Figure 7c)** but had no effect in endothelial cells or keratinocytes **(Figure 7c)**.

**Figure 7:**
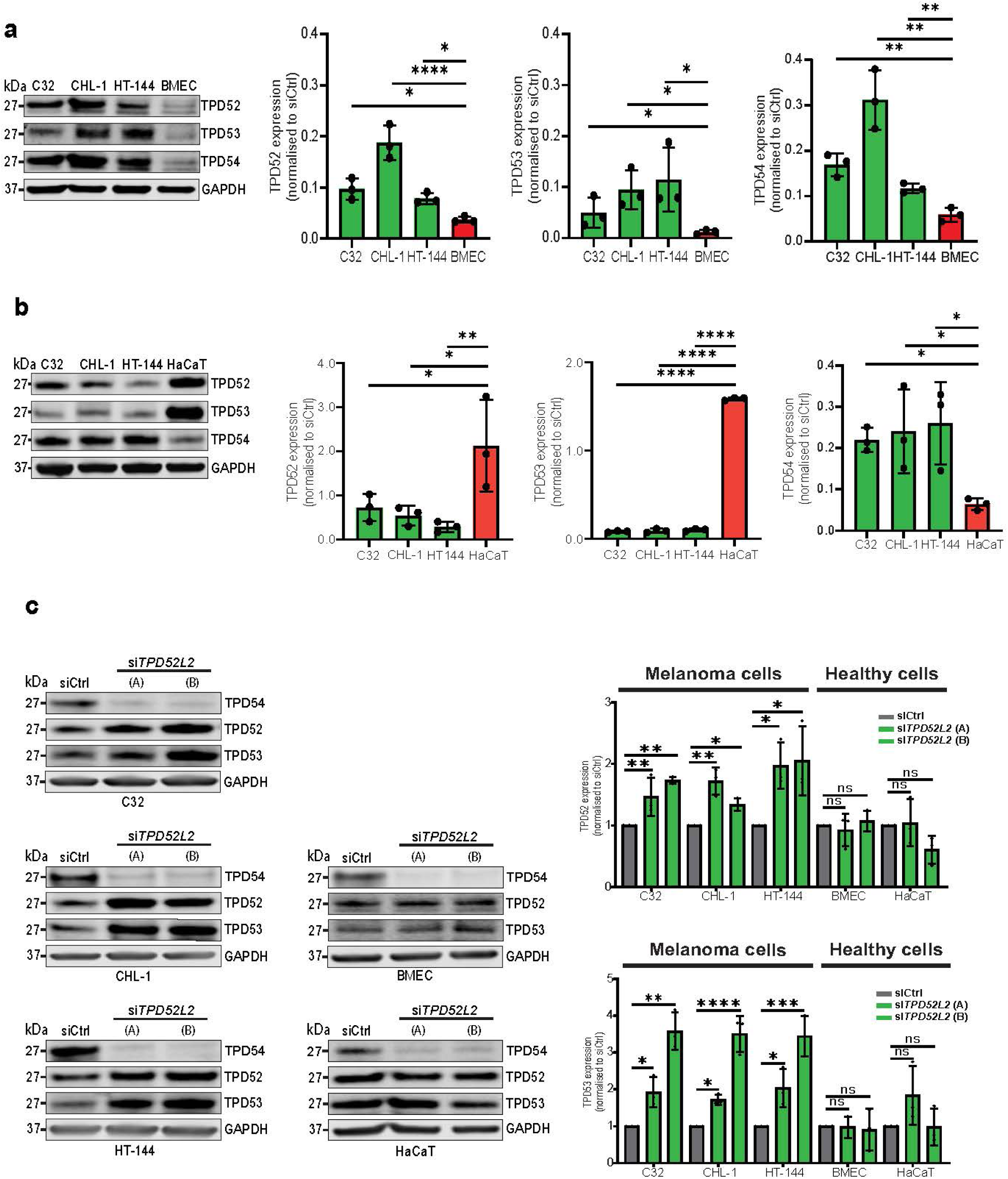
TPD54 expression in melanoma cells, endothelial cells and keratinocytes. **a)** Representative immunoblot and comparative protein expression of TPD52, TPD53 and TPD54 of melanoma cell lysates (C32, CHL-1 and HT-144) versus BMEC endothelial cell lysates (n=3 independent experiments). **b)** Comparative protein expression of TPD52, TPD53 and TPD54 via immunoblot of melanoma cell lysates (C32, CHL-1 and HT-144) versus keratinocytes (HaCaT) cell lysates (n=3 independent experiments). **c)** Protein expression via immunoblots of TPD4, TPD52 and TPD53 in response to *TPD52L2* KD in melanoma cells (C32, CHL-1 and HT-144) versus BMEC endothelial cells and HaCaT keratinocytes. Data are presented as mean ± SEM and analyzed using one-way ANOVA to compare against control groups. * = p<0.05, ** = p<0.01 and **** = p<0.0001

With documentation that TPD54 has ten transcript variants (V1–V10) (66), and that V5 and V6 have high expression across multiple cancers, including breast, lung, and melanoma (66) we examined, for the first time, whether healthy cells (BMEC and HaCaT) might express transcripts other than V5 and V6. This was achieved using the Oxford Nanopore Technologies long-read amplicon sequencing to compare *TPD52L2* within melanoma cells (C32, CHL-1, HT-144), endothelial cells (BMEC), and keratinocytes (HaCaT). Interestingly, only two variants of *TPD52L2* were detected in all cells (cancerous and healthy), V5 and V6 **(Figure 8a)**, and no significant differences in the expression levels of these variants were observed across the healthy and melanoma cell lines **(Figure 8b)**. With additional short-read RNAseq data of different tissues (including PBMCs, skin fibroblasts and urothelial cells) indicating that V5 is the predominant transcript and V6 being the second most abundant (not shown), it is our contention that these are the most relevant transcripts to pursue in cancer biology.

**Figure 8:**
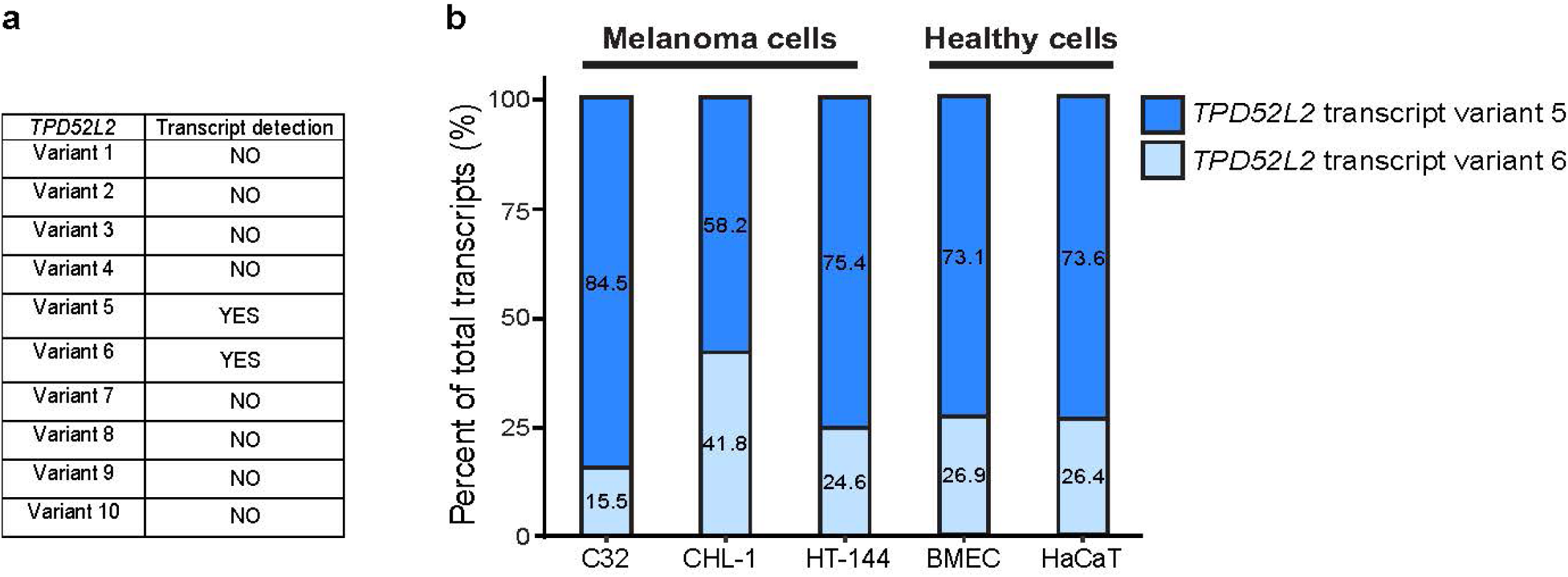
*TPD54* transcript variant expression in melanoma cells, endothelial cells and keratinocytes. **a)** Table of Nanopore long read sequencing of *TPD52L2* transcript variants (V1-V10) in melanoma cells (C32, CHL-1 and HT-144), endothelial cells (BMEC) endothelial cells and keratinocytes (HaCaT). **b)** Percentage of *TPD52L2* variants detected for cells in (**a**).

### Targeting TPD54 reduces melanoma tumor growth and modulates the TME

To investigate whether targeting TPD54 could perturb melanoma growth *in vivo,* we developed a *Tpd52l2* knockout (KO) via *CRISPR-Cas9* editing in the murine melanoma cell lines B16-F10 and HCmel12 **(Figure 9a-d, Sup Figure S3a)**. In keeping with the siRNA analysis of human cells, *in vitro* functional analysis of *Tpd52l2* KO B16-F10 and HCmel12 cells demonstrated no change in proliferation but lower cell migration when compared to non-targeting control (NTC) cells **(Sup Figures S3b-g)**. The B16-F10 and HCmel12 cells (without and with *Tpd54* knockout) were engrafted subcutaneously into both side flanks of C57BL/6 mice and tumor growth was monitored via calipers twice weekly for 12-20 days (ethical endpoint). **Figures 9e-f** show a significantly lower tumor burden in mice engrafted with the *Tpd54* KO tumors when compared to those engrafted with NTC controls. We further took the opportunity to investigate the metastatic potential of B16-F10 cells (NTC or *Tpd52l2* KO) via intravenous administration into C57BL/6 mice, with lungs harvested 14 days post injection. **Figure 9g** shows a significantly lower metastatic burden in the lungs of mice injected with *Tpd52l2* KO B16-F10 cancer cells compared to those injected intravenously with NTC cells.

**Figure 9:**
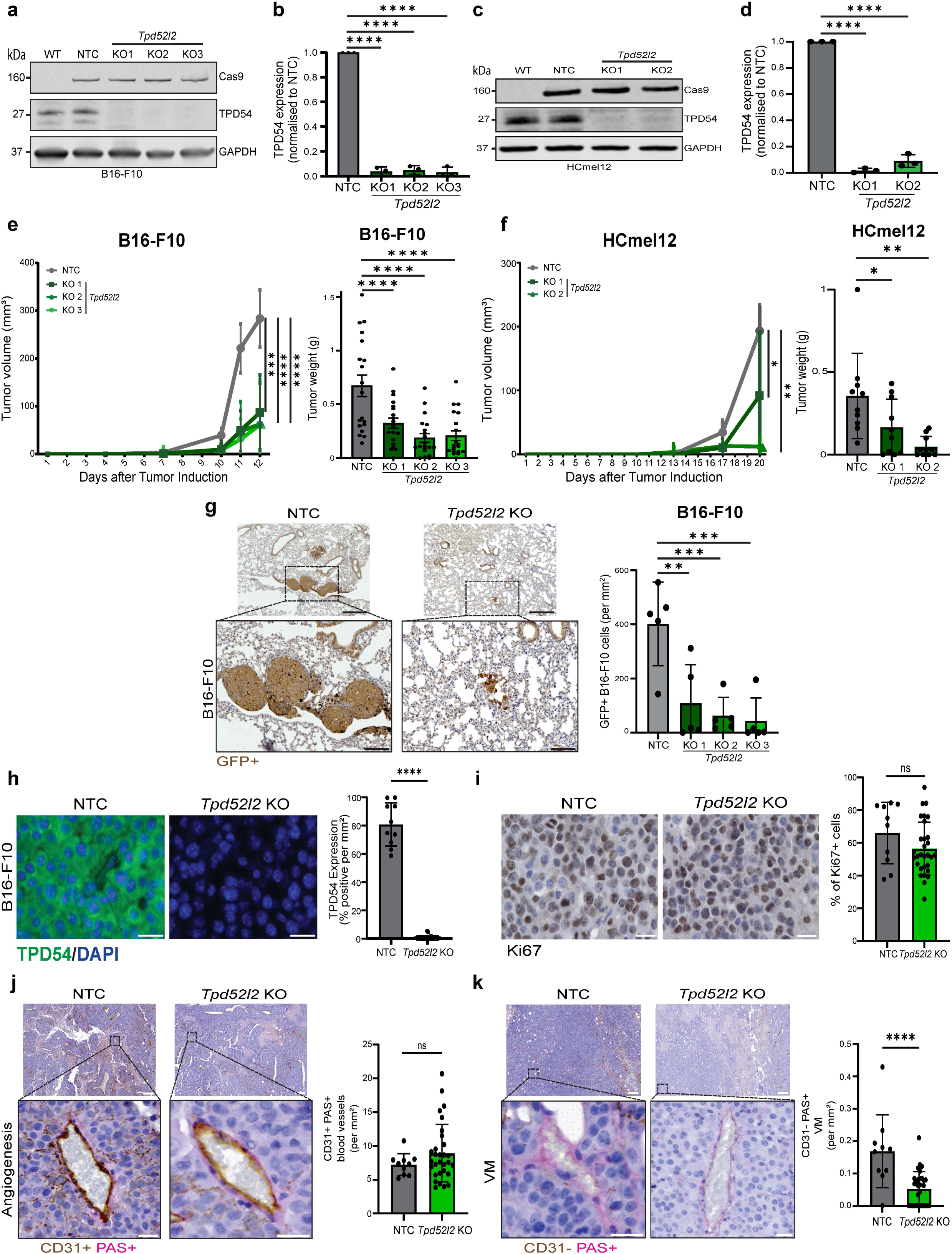
Targeting TPD54 to combat melanoma *in vivo.* **a-d)** Representative immunoblots and quantification of *Tpd52l2* CRISPR-Cas9 knockout (KO) in murine B16-F10 (a-b) and HCmel12 (c-d) melanoma cells (n=3 independent experiments). **e-f)** Caliper measurements and tumor weight at experimental endpoint for B16-F10 (day 12) and HCmel12 (day 20) without and with *Tpd52l2* KO versus non-targeting control (NTC) tumors**. g)** Representative IHC images of GFP staining of B16-F10-GFP-luc melanoma cells in lung tissue of control (NTC) and *Tpd52l2* KO cancer bearing mice at end point (day 12). Quantification of GFP+ melanoma cells per mm^2^ per lung tissue, scale bar = 500 μm (top) and 100 μm (bottom) (n=10-30 tumors per group) **h)** Representative images (left) and quantification (right) of IF staining for TPD54 in NTC and *Tpd52l2* KO B16-F10 tumors at end point (day 12), with TPD54 (green), nucleus (DAPI, blue). Scale bar = 50 μm., (n=10-30 tumors per group). **i)** Representative IHC images of tumor cell proliferation marker Ki-67 in NTC and *Tpd52l2* KO tumors at end point (day 12). Right: quantification of %Ki-67+ cells per mm^2^ per tumor, scale bar = 20 μm, (n=10-30 tumors per group). **j)** Representative IHC image of CD31+ endothelial cell lined blood vessels detected via IHC staining with Periodic-acid Schiff (PAS) stain in B16-F10 NTC and *Tpd52l2* KO tumors at Day 12, scale bar = 200 μm (top) and 20 μm (bottom). Tumor angiogenesis quantified as the number of CD31+/PAS + blood vessels containing blood cells per mm^2^ per tumor (n=10-30 tumors per group). **k)** Representative images of VM detected via IHC staining of CD31-/PAS+ in B16-F10 NTC and *Tpd52l2* KO tumors at endpoint (Day 12). Scale bar = 200 μm (top) and 20 μm (bottom). VM vessel density was quantified as the number of CD31-/PAS+ VM vessels containing blood cells per mm^2^ per tumor. (n=10-30 tumors per group). Data are presented as mean ± SEM and analyzed using one-way ANOVA to compare to NTC groups. ns= >0.05, * = p<0.05 and **** = p<0.0001

Analysis of the B16-F10 tumors established in the flanks of mice confirmed depletion of TPD54 protein in the *Tpd52l2* KO cell line only **(Figure 9h)** but a similar proportion of Ki67+ cell cycling **(Figure 9i)**, and no difference in apoptosis **(Sup Figures 3h, i)**. Given the vital role the TME plays in promoting cancer progression (8, 9, 67, 68), we next examined the vasculature of the B16-F10 tumors and observed no difference in the abundance of CD31+PAS+ vessels **(Figure 9j)**. However, significantly less vasculogenic mimicry (CD31-PAS+ VM vessels) was observed in the *Tpd52l2* KO B16-F10 tumors **(Figure 9k)** compared to controls. Mass cytometric analysis showed that the B16-F10 *Tpd52l2* KO tumors had more CD45⁺ immune cells when compared to the control tumors (**Figure 10a** with gating of live cells and leukocyte subsets shown in Sup Figure S4). More specifically, the B16-F10 *Tpd52l2* KO tumors contained a higher proportion of CD3⁺ T cells, CD8⁺ T cells, and natural killer (NK) cells **(Figure 10a),** but a lower proportion of myeloid cells and dendritic cells when compared to control tumors **(Figure 10a)**. Finally, given the aforementioned observations of TPD54 regulating the melanoma secretome *in vitro*, we examined the circulating chemokines and cytokines via a protein profiler array. **Figure 10b** shows that the pooled serum from six mice harboring B16-F10 *Tpd52l2* KO tumors contained less circulating chemokine CCL6, fibroblast growth factor FGF-21, osteoprotegerin and more insulin growth factor binding protein IGFBP-5 and sE-selectin when compared to mice with B16-F10 NTC tumors.

**Figure 10:**
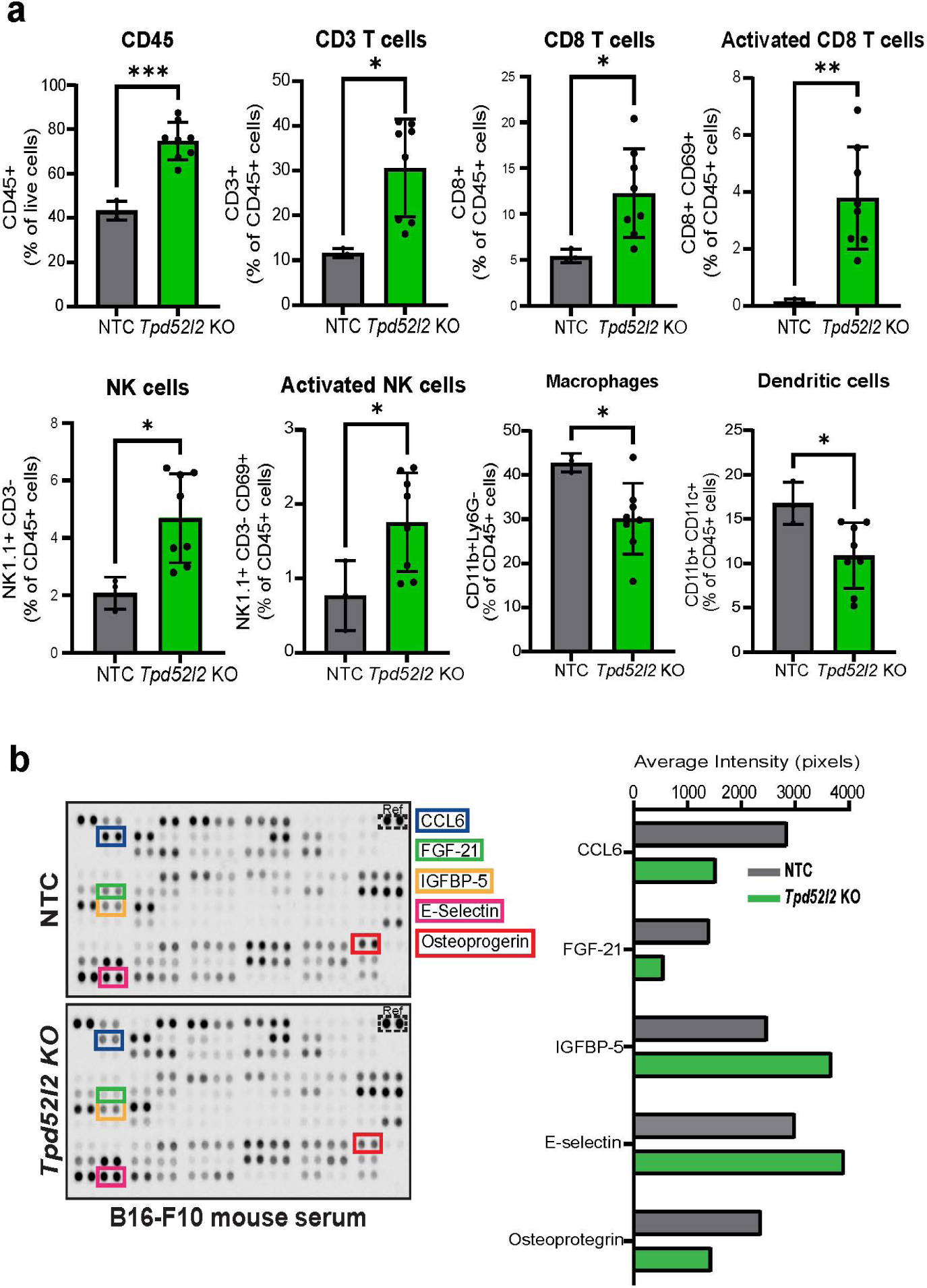
Targeting TPD54 in B16-F10 melanoma cells alters tumor infiltrating leukocytes *in vivo*. **a)** The percentage of CD45+ leukocytes within the live cell population, followed by leukocyte subset identification presented as a percentage of the CD45+ population (n=3-8 tumors per group). Data are presented as mean ± SEM and analyzed by one-way ANOVA. ns= >0.05, * = p<0.05, ** = p<0.01, *** = p<0.001 **b)** Representative images and quantification of proteome profiling cytokine array on pooled sera from mice with B16-F10 tumors (NTC n=6 mice) and (*Tpd52l2* KO n=6 mice). Data shown from pooled samples on a single array experiment.

## Discussion

Cancer progression is not only characterized as a stepwise accumulation of genetic and epigenetic aberrations, but also a resulting pathological rewiring of fundamental cellular processes (69–71). To sustain growth and dissemination, malignant cells must continuously produce and distribute pro-tumorigenic molecules, remodel intracellular organelles, and secrete a diverse repertoire of modulatory factors that collectively promote invasion, vascularization, and immune evasion (17, 44, 57, 72–75). Central to this malignant adaptability is the membrane trafficking system, which integrates the endoplasmic reticulum, Golgi apparatus, endosomal dynamics and EV secretion to coordinate both tumor-intrinsic plasticity and TME remodeling (17, 44, 57, 72–75).

In this study, we identify the intracellular transport protein TPD54 as an underappreciated component of melanoma progression, providing insights into both its biological functions and implications for patient outcomes. We show that *TPD52L2* expression is selectively upregulated in melanoma, correlating with poor survival and reduced responsiveness to immune checkpoint blockade and targeted therapy. This clinical association is consistent with the established paradigm that aberrant gene expression programs underpin tumorigenesis and the development of therapeutic resistance (76). Mechanistically, in melanoma, TPD54 preserves Golgi integrity, sustains Rab GTPase dependent trafficking, and mediates the cell surface expression (and recycling) of adhesion-signaling molecules such as integrin-β1 and DSG2. Collectively, these functions enable melanoma cells to maintain dynamic cycles of membrane remodeling that are indispensable for cancer cell signaling, adhesion, migration, and invasion (17, 44, 48, 49, 52, 55, 57, 72–75, 77, 78). The disruption of this pro-cancerous program following TPD54 depletion highlights the reliance of melanoma cells on intact vesicular transport to sustain their cellular homeostasis and malignant function.

Beyond intracellular trafficking, we established TPD54 as a regulator of the melanoma secretome through its modulation of EV release and cytokine secretion. Tumor-derived EVs are increasingly recognized as essential messengers that extend the influence of malignant cells beyond their immediate niche, disseminating oncogenic proteins, nucleic acids, and immunosuppressive mediators to distant stromal, vascular, and immune compartments (15, 16, 18, 79–81). Depletion of TPD54 not only markedly reduces EV output but also altered the secretion of angiogenic (e.g., DKK1) and immunomodulatory factors (e.g., MCP-1, IL-8, TNF-α) in melanoma cells. These findings position TPD54 as a molecular coordinator that couples intracellular cargo recycling with EV secretion to fine-tune melanoma cell functionality.

A particularly striking outcome of TPD54 activity in melanoma was its essential role in vasculogenic mimicry (VM). Unlike endothelial cell mediated angiogenesis, VM reflects the remarkable plasticity of melanoma cells, enabling them to form perfuseable, vessel-like structures that support nutrient delivery and metastatic dissemination (49, 60, 61, 82–84). We observed that TPD54 is required for VM in melanoma cells but is dispensable for endothelial cell angiogenesis, suggesting a cancer cell-specific regulatory mechanism. This selectivity is consistent with the higher expression of TPD54 in melanoma compared to endothelial cells (and keratinocytes) as well as our observation that targeting TPD54 in endothelial cells did not impact protein trafficking as was seen in melanoma cells (e.g Golgi integrity and DSG2 trafficking). This disparity may reflect the disproportionate vesicular trafficking demands of malignant cells compared to healthy cells, driven by the need to produce and deliver high levels of proteins that facilitate cancer-associated processes, including migration (8, 15, 17, 57, 73, 75, 80, 85, 86).

Beyond the cancer cell autonomous role, our study uncovered a pivotal role for TPD54 in modulating melanoma tumor development via the TME, particularly the vasculature and immune cell landscape. Loss of TPD54 in murine melanoma models caused a significant reduction in tumor burden, metastasis, impaired VM, as well as a pronounced reprogramming of the immune landscape. More specifically, inhibition of *Tpd52l2* enhanced the infiltration of CD8⁺ T cells and NK cells, coupled with a reduction in macrophages and dendritic cells. This shift toward a more immunostimulatory TME suggests that elevated TPD54 in cancer contributes to immune evasion, likely through modulating the secretion of cytokines and other immunosuppressive mediators. These observations are consistent with published *in silico* gene enrichment analyses, which suggest that *TPD54* expression is positively associated with regulatory T cells (Tregs), tumor-associated macrophages (TAMs), and immunosuppressive genes such as TGFβ1, while being negatively correlated with CD8⁺ T cells and NK cells (34).

An unexpected observation was that while inhibition of TPD54 prevented VM formation (*in vitro* and *in vivo*) it had no effect on angiogenesis by endothelial cells. Knockdown of TPD54 in endothelial cells and keratinocytes also had no effect on Golgi integrity or the transport of proteins to the plasma membrane, thus raising the possibility that an oncogenic form of TPD54 exists. With long read sequencing identifying the same genomic transcripts of TPD54 in both cancerous and healthy cells (i.e. variants 5 and 6 at ∼75% and ∼25%, respectively), the molecular mechanisms underpinning an ‘oncogenic’ form of TPD54 is yet to be elucidated. If a cancer specific TPD54 can be identified, it is tempting to speculate that it could potentially be exploited as a therapeutic target to selectively disrupt tumor-derived vascular networks while sparing physiological vasculature and healthy cell function. Additional studies are needed to substantiate and extend these observations.

Together, our findings position TPD54 at the intersections of vesicular trafficking, protein transport, tumor growth, and immune cell evasion, revealing it to be more than just a carrier protein and perhaps more of a ‘control node’ that couples processes extending from the transport of intracellular cargo to extracellular immune programming. By orchestrating the recycling of adhesion proteins, release of EVs, and cytokine secretion, it is tempting to speculate that TPD54 integrates melanoma cell migration, VM, and immune suppression into a unified trafficking-centered axis for cancer progression. This central role highlights TPD54 as a previously unrecognized regulator of melanoma adaptation, offering critical insight into the molecular mechanisms that underpin disease progression. The association of TPD54 with immunotherapy resistance further raises the possibility that targeting trafficking regulators could synergize with immune checkpoint inhibitors by restoring effector immune function and dismantling tumor-derived immunosuppressive networks (87). Thus, TPD54 emerges as an underappreciated regulator of melanoma with a translational potential that warrants further investigation to delineate the molecular interactome of TPD54 and to evaluate TPD54-targeted strategies in preclinical models of melanoma and beyond.

## Methods

### Ethical Statements

#### Human ethics

The use of archived tumor tissue sections and matched clinical data from melanoma patients were given ethical clearance by the Central Adelaide Local Health Network (CALHN). Ethics approval and patient consent granted under (HREC/16/RAH/95). Deidentified tumor microarray (TMA) sections and matched clinical data were purchased via the TissueArray BioResource biobank (cat # ME204, Derwood, MD, USA,).

#### Animal ethics

Animal experiments were given ethical clearance from the University of South Australia Animal Ethics Committee in compliance with the Australian Code for the Care and Use of Animals for Scientific purposes, with assigned project number U11-23.

### Cell culture

The human melanoma cell line C32 was purchased from CellBank Australia (Westmead, NSW, Australia), human melanoma cell lines CHL-1 and HT-144 were gifted from Prof. G McArthur (Peter MacCallum Cancer Centre, Melbourne, Vic, Australia) and keratinocytes (HaCaT) were cultured as published (49). Trabecular human bone marrow endothelial cell line (TrHBMEC (BMEC)), gifted from B Weksler (Cornell University Medical College, New York, NY, USA)(64), enriched for DSG2+ cells and cultured as published (65, 88, 89). All human cell line’s identity was confirmed by PCR-based short tandem repeat (STR) analysis (SA Pathology, Adelaide, SA, Australia). Mouse melanoma cell line B16-F10-GFP-P2A-luciferase (referred to as B16-F10) was gifted from Prof. Jeff Holst (University of Sydney, Sydney, NSW, Australia) and maintained in DMEM (Gibco) with 10% FBS (HyClone). HCmel12 was gifted from Prof. T Bald (University of Queensland, Brisbane, Qld, Australia) and cultured as published (90). All cells were maintained in a humidified 5% CO_2_ incubator at 37°C. Cultures were routinely checked for mycoplasma absence via MycoAlert (Lonza, Basel, Switzerland).

### *In silico* gene analysis

*TPD52, TPD53, TPD54* expression plots and Kaplan–Meier survival analysis using GEPIA2 (35) downloaded from website http://gepia2.cancer-pku.cn/#index Kaplan–Meier survival analysis of melanoma patients treated with immune therapy was performed using the KMPlotter tool (37): https://kmplot.com/analysis/index.php?p=service&cancer=immunotherapy Analysis of *TPD52*, *TPD53* and *TPD54* relevance in human melanoma disease stage and BRAF+MEK inhibitor treatment was downloaded from NCBI and analyzed using *GEO2R analysis tool* http://www.ncbi.nlm.nih.gov/geo/geo2r., datasets are available under accession code GDS1965, GSE196434. Melanoma patient response to immune therapy were downloaded from https://rocplot.org/immune (91).

### Small interfering RNA transient knock down of *hTPD52L2*

‘Trilencer-27’ siRNA duplexes were purchased from Origene (Rockville, MD, USA). Transfection was performed using Lipofectamine RNAiMax and OptiMEM reduced serum media (Life Technologies, Thermo Fisher Scientific, Waltham, MA, USA) according to the manufacturer’s protocol. Cells were incubated with the duplex master-mix for 72hrs. Control cells were treated with equivalent amounts (10nM) of the universal non-silencing siRNA duplex (Origene). Knockdown efficiency was routinely checked using immunoblotting.

### CRISPR/Cas9 knock out of *Tpd52l2*

To knockout *Tpd52l2* in the murine B16-F10 and HCmel12 cells, HEK293T cells were first transfected with a master mix containing OptiMEM reduced serum media (Life Technologies), Lipofectamine 2000 (Invitrogen, Carlsbad, CA, USA), 5µg of DNA for each lentiviral packaging construct; pLP1, pLP2, pTAT and pVSVG (ViraPower Lentiviral systems kit, Invitrogen), and 15µg of DNA for either a CRISPR non-targeting sgRNA All-in-One-Lentiviral Vector (with spCas9) or the *Tpd52l2* sgRNA CRISPR-Cas9 All-in-One Lentivector Set (mouse, Applied Biological Materials, Richmond, Canada). Following 24 hrs of incubation, the medium was replaced with cell specific media. Viral supernatant was collected after a further 48 hrs of incubation, centrifuged at 845 g for 5 mins at 4°C, and filtered using a 0.45 µm filter. Freshly harvested virus was mixed with media containing 8µg/mL polybrene (Merck, Darmstadt, Germany) before being added to target cells for transduction. The transfected cells underwent a two-week period of puromycin selection at 2 µg/mL. To obtain individual clones for B16-F10, cells were single cell sorted using flow cytometry on the FACSAria™ Fusion Flow Cytometer (BD Biosciences, NJ, USA), with knockout efficiency assessed through immunoblotting. The CRISPR/Cas9 knockout targets used to silence Tpd54 had the following sequences: Non-targeting control NTC: ATCTCGCTTGGGCGAGAGTAAG; *Tpd52l2* KO.1: TCAAAGACGAATATGCCGAA; *Tpd52l2* KO.2: CATCACTAGCATTCTTGACC; *Tpd52l2* KO.3:ACTTTGATGCGTAGTTCCAA.

### Immunocytochemistry

Cells were seeded onto coverslips and fixed by 4% paraformaldehyde (PFA) (VWR International, PA, USA) for 10min and permeabilized by 0.25% PBST (Triton X-100 (Merck) in PBS) 10min. Cells were washed with cold PBS and then blocked by normal goat serum (1:10) in 3% BSA/PBS for 3hr and then incubated with primary antibodies overnight at 4°C. The following day, the coverslips were washed with PBS and incubated with a species-specific Alexa-Fluor secondary antibody (Thermo Fisher) and DAPI (Merck) for nuclear staining for 1 hr at RT in the dark. The coverslips were washed with PBS and mounted onto glass slides using 10 μL Fluoro-Gel water based mounting medium (ProSciTech, QLD, Australia) and cured for 24 hrs at RT in the dark. Immunofluorescence images were captured on the LSM 800 Confocal Microscope (Zeiss, Jena, Germany) or Axioscan 7 (Zeiss) and processed using the Zen 2011 software (Zeiss) and ImageJ. Primary antibodies used anti-TPD54 (United Bioresearch, Salt Lake City, UT, USA), anti GM130 (BD Bioscience, San Jose, CA, USA), anti-TGN46 (Abcam, Cambridge, UK), anti-integrin-β1 (BD Bioscience), anti-DSG2 (R&D Systems, Minneapolis, MN, USA), anti-Rab5, anti-Rab4, anti-Rab7, anti-Rab9, anti-Rab10 (all from Cell Signaling Technology, Danvers, MA, USA).

### Histology, tissue processing and sample preparations

Tumor tissue was dissected, treated with 4% PFA overnight, processed, and formalin-fixed, and paraffin embedded (FFPE). Tissue blocks were then cut into 4-5μm sections and mounted onto glass slides for histological analysis. Heat induced antigen retrieval was undertaken in citric acid (pH 6.5) buffer. Sections were then subjected to staining via immunohistochemistry (IHC) or immunofluorescence (IF) labeling.

#### Immunohistochemistry

VECTASTAIN elite ABC HRP/Peroxidase kit: Following antigen retrieval, sections were incubated with 3% hydrogen peroxide for 10min and blocked with 5% normal serum diluted in 1x PBS for 1 hr, followed by an overnight incubation with primary antibodies diluted with 5% normal serum in 1x PBS in a humidified chamber. The following day, sections were incubated with biotinylated secondary antibodies (1:500) for 35min at RT, followed by processing with a VECTASTAIN elite ABC HRP/Peroxidase kit (Vector Laboratories, Newark, CA, USA) for 30 min and visualized with a DAB+ substrate kit (Vector Laboratories), as per manufacturer’s instructions. Sections were then counter stained with hematoxylin to visualize nuclei (Agilent Technologies, Santa Clara, CA, USA), dehydrated, and mounted in Dibutylphthalate Polystyrene Xylene (DPX) medium (Merck).

DAKO REAL EnVision detection system: Following antigen retrieval, sections were incubated with 0.5% hydrogen peroxide diluted in methanol for 10min and blocked with 5% normal serum diluted in 1x TBS for 1hr at room temperature, followed by an overnight incubation at 4°C with primary antibodies diluted with 3% normal serum in 1x TBS in a humidified chamber. The following day, sections were incubated with the DAKO envision plus dual link system HRP for 2.5hr at RT in a humidified chamber, as per the manufacturer’s instructions. Following incubation, slides were washed three times in 1x TBS. Slides were incubated with the DAKO liquid DAB+ substrate chromogen cocktail to visualize staining, as per manufacturer’s instructions. Sections were then counter stained with hematoxylin to visualize nuclei (Agilent Technologies), dehydrated, and mounted in Dibutylphthalate Polystyrene Xylene (DPX) medium (Merck). Images captured via a Hamamatsu Nanozoomer NDP slide scanner (Hamamatsu Photonics, Shizuoka, Japan), visualized using NDP Virtual Slide Viewer, and quantification performed using QuPath Digital Pathology Image Analysis software (University of Edinburgh, Edinburgh, UK) or ImageJ.

#### Immunofluorescence

Following antigen retrieval, slides were blocked with 5% normal serum in casein block, CAS-Block™ (Thermo Fisher Scientific) for 30-45min, followed by primary antibody incubation overnight in 4°C and a secondary antibody incubation the subsequent day for 1hr at RT. All antibody dilutions were made up in CAS-Block™ with 5% normal serum. Slides were mounted in a Fluoro-Gel water based mounting medium (ProSciTech) and left to cure for 24 hr at RT in the dark. Imaging of sections was performed using an LSM 800 confocal microscope (Zeiss) or Axioscan 7 (Zeiss) and processed using the Zen 2011 software (Zeiss) or ImageJ Fiji (NIH).

### Extracellular vesicle harvest

Harvested cell conditioned media was cleared of cells and cell debris by sequential centrifugations at 800g for 10min followed by 2,500g for 15min. Cell media underwent ultracentrifugation in the Optima MAX-XP ultracentrifuge (TLA110 rotor, Beckman Coulter, Brea, CA, USA) at 100,000g for 2hr to isolate extracellular vesicles. Supernatant was discarded and the pellet was resuspended in either 40 µl of RIPA buffer for immunoblot analysis or 1 mL of filtered PBS for nanoparticle tracking analysis on the NanoSight NS300 (Malvern Panalytical, Worcestershire, UK).

### Immunoblotting

Cells were lysed directly in dishes with cold RIPA lysis buffer (150 mM NaCl, 50 mM Tris-Cl, 1% Triton, 1% deoxycholate, 0.1% SDS, 2 mM EDTA) supplemented with complete Protease Inhibitor (Sigma-Aldrich). Lysates were collected and clarified. Protein concentration was determined using a Pierce™ BCA protein assay kit (Thermo Fisher) and measured on a FLUOstar Omega Microplate Reader (BMG Labtech, Ortenberg, Germany). Proteins were separated on Criterion XT Bis-Tris Gels (Bio-Rad Laboratories, Hercules, CA, USA) and transferred on nitrocellulose membrane (Pall Corporation, Port Washington, NY, USA), which was then blocked in Tris-buffered saline with Tween-20 (TBST) containing 5% BSA for 3 hr at RT. The membranes were then probed overnight at 4°C with primary antibodies diluted in TBST containing 5% BSA. Primary antibodies used were: TPD52 (1:800, Abcam, ab182578); TPD52L1/TPD53 (1:800, United Bioresearch, 14732-1-AP); TPD52L2/TPD54 (1:800, United Bioresearch, 11795-1-AP; or Thermo Fisher Scientific, PA5-112772); DSG2 (1:4000, Bethyl Laboratories, A303-758A); Cas9 (1:1000, New England Biolabs, 14697S); β-actin (1:1000, Sigma-Aldrich, MAB150); α-tubulin (1:2000, Thermo Fisher Scientific, A11126); GM130 (1:1000, BD Biosciences, 610823); CD81 (1:1000, Cell Signaling Technology, 56039); ALIX (1:1000, Cell Signaling Technology, 2171S); GAPDH (1:3000, Cell Signaling Technology, 5174); and rabbit IgG control (5 µg, Cell Signaling Technology, 2729S). Next, membranes were incubated with IRDye-conjugated secondary antibodies (anti-rabbit or anti-mouse, LI-COR Biosciences) diluted 1:10,000 and signals were visualized and quantified using the Odyssey imaging system and Image Studio software (LI-COR Biosciences, Lincoln, NE, USA).

### Immunoprecipitation

Cells were lysed in lysis buffer (150mM NaCl, 50mM Tris-Cl, 1% Triton, 1% deoxycholate, 0.1% SDS, 2mM EDTA and complete protease inhibitor cocktail (Roche, Basel, Switzerland)), followed by homogenization and centrifugation (13,000 rpm for 8 min at 4°C). Total lysates were incubated with 100µl of Protein G Dynabeads (Thermo Fisher) for 4hr on a rotation wheel, then centrifuged at 13,000rpm for 5min. Lysates supernatant were then incubated with 5µg anti-TPD54 (United BioResearch) and isotype control with anti-IgG (Cell Signaling Technologies) overnight on rotation. 100µl of Protein G Dynabeads was added to lysates and incubated for 4hr at 4°C on rotation. Samples were washed three times with 1 mL of lysis buffer, then centrifuged at 13,000rpm for 5min. Supernatant was discarded and 40 µl per sample of SDS sample buffer was added to elute precipitated proteins. Enriched samples were then subjected to SDS–PAGE and western blotting.

### Chemokine profiling array

The chemokine profile of melanoma cell line HT-144 was assessed using the Proteome Profiler Human Chemokine Array Kit (R&D Systems) according to manufacturer’s instructions with 1 mL of melanoma cell culture supernatant. The membrane was visualized using the ChemiDoc imaging system (BIO-RAD). Quantification of the dots on the membrane was performed using ImageQuant™ TL (GE Healthcare Life Sciences, Chicago, IL, USA).

### Nanopore sequencing

Genomic DNA was extracted and amplified using PCR, with optimized conditions for each sample. For sequencing, a ligation-based library preparation method was employed, where equimolar PCR products were ligated with individual barcodes and sequencing adapters from the Oxford Nanopore Technologies Native Barcoding kit 24 V14 (SQK-NBD114.24, Oxford, UK), and sequenced using the Oxford Nanopore MinION Mk1B. 50fmol of prepared library was loaded onto an Oxford Nanopore MinION flow cell (R10.4.1), and sequencing was performed following the manufacturer’s guidelines. The raw signal data was processed using the Oxford Nanopore base calling software (Guppy), which converted the signals into nucleotide sequences. These sequences underwent quality control procedures to assess metrics such as read length, base accuracy, and overall quality, with low-quality reads discarded. For multiplexed samples, the sequences were de-multiplexed based on their unique barcodes, allowing for the separation of individual samples. The sequencing data were further analyzed for read depth, alignment to reference genomes. The final sequencing data were analyzed using bioinformatics tools such as NanoPlot (92), EPI2ME ((https://github.com/epi2me-labs)), and custom scripts, generating visualizations and quality metrics of read length distributions and base accuracy. The final output consisted of high-quality, de-multiplexed DNA sequences, which were used for variant detection.

### Flow cytometry

Cells were incubated with human IgG (Invitrogen) block for 10 min, prior to the addition of IgG1 isotype (BD Biosciences) or anti-human DSG2 (6D8, Thermo Fisher) diluted in HUVE media (complete Medium199 media (Sigma-Aldrich), 20% FBS (Hyclone)), culture flasks pre-coated with gelatin. Following two washes, 5 µg of anti-mouse-DyLight 650 and 0.25 µg/test viability dye 7-AAD (BD Biosciences) was added for 30 min at 4°C. Washed cells were then resuspended in FACS fix (1% Formaldehyde, 20 g/L glucose, 5 mM sodium azide) and analyzed on the Accuri C6 flow cytometer (BD Biosciences) prior to analysis via FCS Express 7 (De Novo Software, Pasadena, CA, USA).

DSG2 resurfacing experiments used melanoma cells (at 90% confluence), placed on ice, treated with 500 µL of cold 0.3M glycine/1% BSA (acid wash) for 10min prior to neutralization with 1ml of complete RPMI media. Cells were returned to the 37°C CO_2_ incubator for 6hr prior to DSG2 assessment via the flow cytometry protocol above.

### Angiogenesis and Vasculogenic Mimicry assays

Cancer cells or endothelial cells were added in wells of μ-angiogenesis culture slide (Ibidi, Grafeling, Germany) on top of growth factor reduced (GFR) basement membrane matrix Matrigel (Invitrogen) then incubated at 37°C (humidified 5% CO2) for 12-24hr under ambient oxygen conditions. Images of the Matrigel assay were captured using an inverted imaging microscopy (EVOS XL).

### Cell proliferation assay

Proliferative activity was assessed over 2 hr at 37°C using the alamarBlue fluorescent dye assay (Invitrogen) as per manufacturer instructions, with fluorescence intensity measured (530-nm excitation and 595-nm emission) using a FLUOstar Optima plate reader (BMG Labtech).

### Cell migration assay

Melanoma cells were harvested and seeded into 8.0μm pore Transwell polycarbonate membrane cell culture inserts (Nunc, Thermo Fisher Scientific) with a chemotactic gradient of 0% (top chamber) to 10% (bottom chamber) serum containing medium. Cells were allowed to transmigrate for 8-24hr in humidified 5% CO2 at 37°C. Transmigrated melanoma cells on the bottom side of the Transwell insert were then visualized by fixing the insert in 4% PFA for 10min at RT and staining the inserts for 5mins with Crystal Violet dye. The inserts were imaged on an inverted imaging microscope (EVOS XL, Life Technologies), and migration index analyzed by detecting stained cells using Image J software (1.47v).

### Cell survival assay

Cell survival was determined by staining with Annexin V-APC (BD Bioscience) for 15min at RT followed by washing and 0.25ug/test 7-AAD (BD Biosciences) staining for 5min at RT. Washed cells were resuspended in FACS fix and analyzed on the Accuri C6 flow cytometer as above.

### Syngeneic subcutaneous melanoma mouse model

Male and female C57Bl/6 mice (6-10 weeks of age) were bred in-house or purchased from Australian Bio Resources (ABR, Garvan Molecular Genetics, Sydney, NSW, Australia) and housed under specific pathogen-free conditions with a standard rodent diet. B16-F10-GFP-P2A-luciferase (B16-F10) cells (1×10^6^) or HCmel12 (2×10^5^) were resuspended in 100μl PBS and injected subcutaneously into both flanks of anaesthetized mice as per ethics approval. Tumor growth was monitored twice weekly via caliper measurements, whereby width and length of tumors were measured to calculate tumor volume based on a modified ellipsoid (tumor volume = ½ (length x width^2^)) (93).

### B16-F10 metastasis mouse model

B16-F10-GFP-P2A-luciferase (B16-F10) melanoma cells (3 × 10⁵ cells) were resuspended in 100μl sterile PBS and administered via tail vein injection into anaesthetized C57Bl/6 mice (male and female (6–10 weeks of age)). Mice were monitored daily for signs of distress and humanely killed 14 days post-injection. Lungs were harvested, fixed in formalin, and processed for paraffin embedding. Pulmonary metastases were evaluated by IHC using an anti-GFP antibody (1:100**, A**bcam, ab6673) and quantified by assessing GFP-positive tumor cells within lung sections via the QPath analysis software.

### Tumor dissociation and CyTOF mass cytometry

Tumor tissues were dissected and dissociated using a mouse dissociation kit (Miltenyi Biotec, Bergisch Gladbach, Germany), using a protocol involving the addition of specific enzymes and utilizing the gentleMACS dissociator (Miltenyi Biotec). Freshly isolated total cell preparations were filtered through 70 μm filter (Falcon, Corning Life Sciences, Tewksbury, MA, USA) and lysed of red blood cells with RBC lysis buffer (0.15M NH_4_Cl, 10mM KHCO_3_).

Cells isolated from mouse tumors were stained for viability in PBS containing 5mM Cell-ID Cisplatin-195Pt (Fluidigm, San Francisco, CA, USA). Cells were then incubated with Human TruStain FcX™ (Fc-Receptor Blocking Solution) (Biolegend, San Diego, CA, USA) for 10min at RT, followed by staining with metal conjugated antibodies for cell surface molecules (Maxpar Mouse Spleen/Lymph node Phenotyping Panel Kit; Fluidigm; 201306) according to the manufacturer’s protocol. Cells were suspended in Maxpar water supplemented with 10% EQ Four Element Calibration Beads (Fluidigm; 201078) and analyzed on a Helios instrument (Fluidigm) with Cytobank premium software. Data was further analyzed using FCS Express 7 Flow Cytometry: Research Edition (De Novo Software).

### Statistics

All statistical analyses and figure generation were performed using GraphPad Prism 10 (GraphPad Software, San Diego, California, USA), except for percent total transcript analysis and figure generation of TPD54 Nanopore sequencing which was performed using R version 4.4.1. Survival analysis was conducted using Kaplan-Meier plots with log-rank tests. Statistical analyses were performed using one-way repeated measures when comparing between samples or two-tailed unpaired Student’s t-tests. Results with p<0.05 were considered statistically significant, with *= p<0.05, **= p<0.01, ***=p<0.001 and ****=p<0.0001 indicated on graphs and figure legends.

## Acknowledgements

This work was funded by a grant to CSB from NHMRC (Ideas Grant GNT2021009). MO was supported by an RTP/USAPA post-graduate scholarship and a Top-Up Scholarship from Tour de Cure. We thank Samantha Escarbe for her expertise in histology and Ian Dempster for sharing his lived experience of metastatic melanoma. BioRender was used to generate graphical schematics.

