## Supplementary Data for "Tumor Protein D54 (TPD54) regulates intracellular protein trafficking, cellular function and disease progression in melanoma"

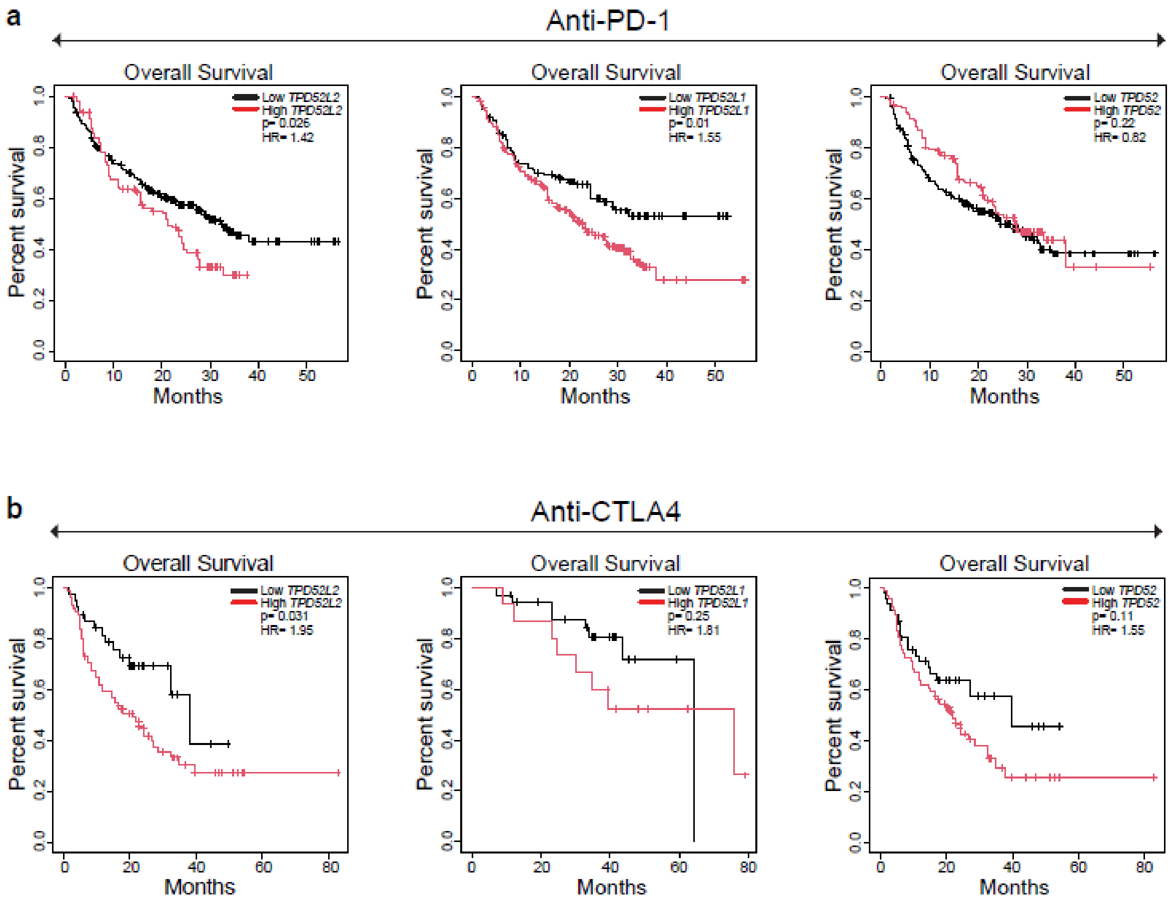


**Supplementary Figure S1: TPD52 family expression relative to patient survival on immune therapy.** **a)** Kaplan-Meier plot from The Kaplan-Meier plotter with RNA sequencing data from the TCGA, Geo and EGA database comparing survival between patients with *TPD52L2, TPD52L1 or TPD52*-high expressing tumors (n=119-227) and *TPD52L2, TPD52L1 or TPD52*-low expressing tumors (n=107-215) which have been treated with PD-1 immune therapy (Nivolumab and Pembrolizumab), log-rank p=0.22, p=0.01 and p=0.026 respectively. **b)** Kaplan-Meier plot from The Kaplan-Meier plotter with RNA sequencing data from the TCGA, Geo and EGA database comparing survival between patients with *TPD52L2, TPD52L1 or TPD52*-high expressing tumors (n=34-46) and *TPD52L2, TPD52L1 or TPD52*-low expressing tumors (n=15-74) which have been treated with CTLA-4 immune therapy (Ipilimumab), log-rank p=0.11, p=0.25 and p=0.031 respectively.

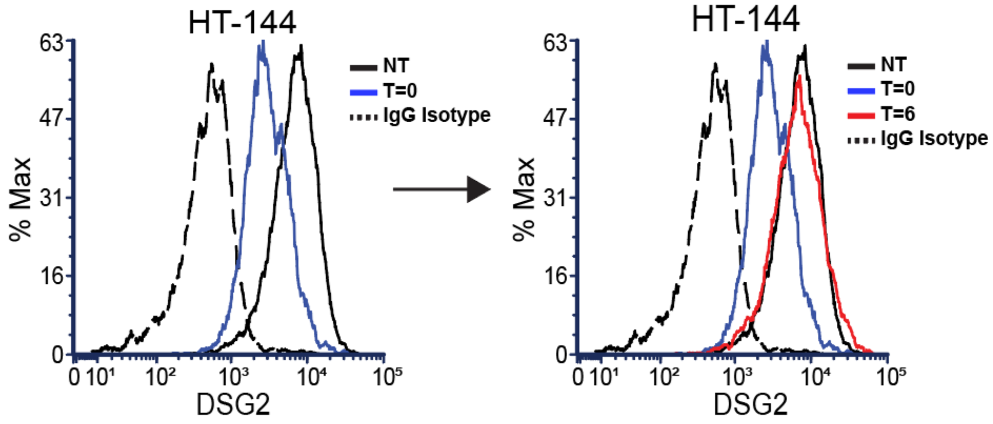


**Supplementary Figure S2: Recycling of DSG2 to the plasma membrane of melanoma cells.**
Exemplar histogram of DSG2 resurfacing via acid buffer treatment on HT-144 melanoma cells. T=0 (time point 0hr), T=6 (time point 6hr).

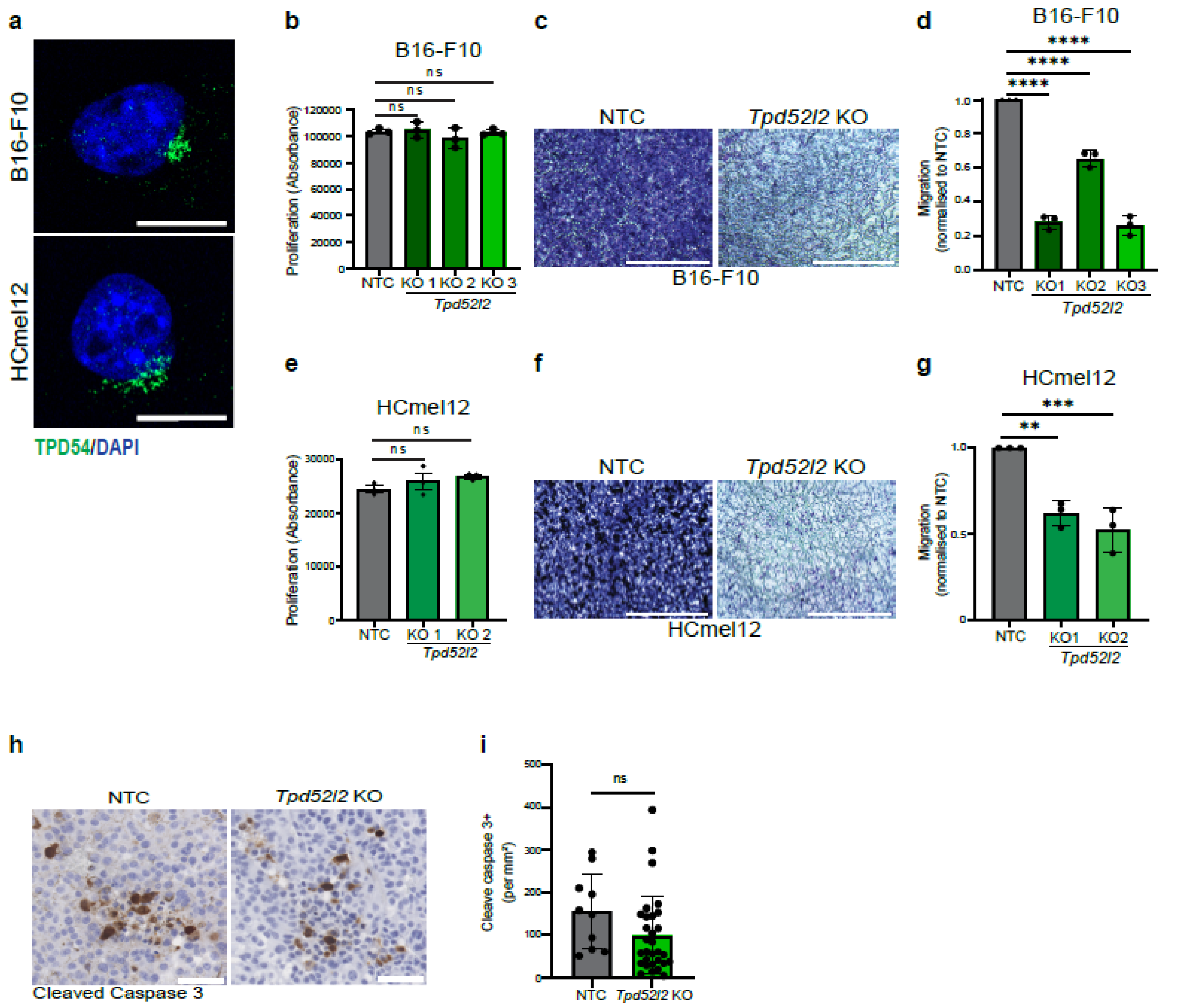


**Supplementary Figure S3: TPD54 expression in mouse melanoma cells, proliferation, migration and survival in tumor.**

**a)** IF images of TPD54 (green) in mouse melanoma cell lines B16-F10 and HCmel12. **b)** AlamarBlue assay to assess cell proliferation capabilities by B16-F10 *Tpd52l2* KO versus NTC cells. **c)** Migration of B16-F10 cells (± *Tpd52l2* KO) in transwell chemotactic assays. Scale bar = 1000µm**. d)** Quantification of migrated cells from c), via crystal violet staining and ImageJ analysis. n=3 independent experiments. **e)** AlamarBlue assay to assess cell proliferation capabilities by HCmel12 *Tpd52l2* KO versus NTC cells. **f)** Migration of HCmel12 cells (± *Tpd52l2* KO) in transwell chemotactic assays. Scale bar = 1000µm**. g)** Quantification of migrated cells from c), via crystal violet staining and ImageJ analysis. n=3 independent experiments. **h)** Representative images of IHC staining of tumor cell apoptosis marker cleave caspase-3 in HCmel12 (± *Tpd52l2* KO) tumors at Day 14. **i)** quantified for cleave caspase 3+ cells per mm$^2$ of the whole tumor section. All graphed data are presented as mean ± SEM and analyzed using one-way ANOVA to compare against control groups. *=p<0.05, **=p<0.01, ***=p<0.001 and ****=p<0.0001

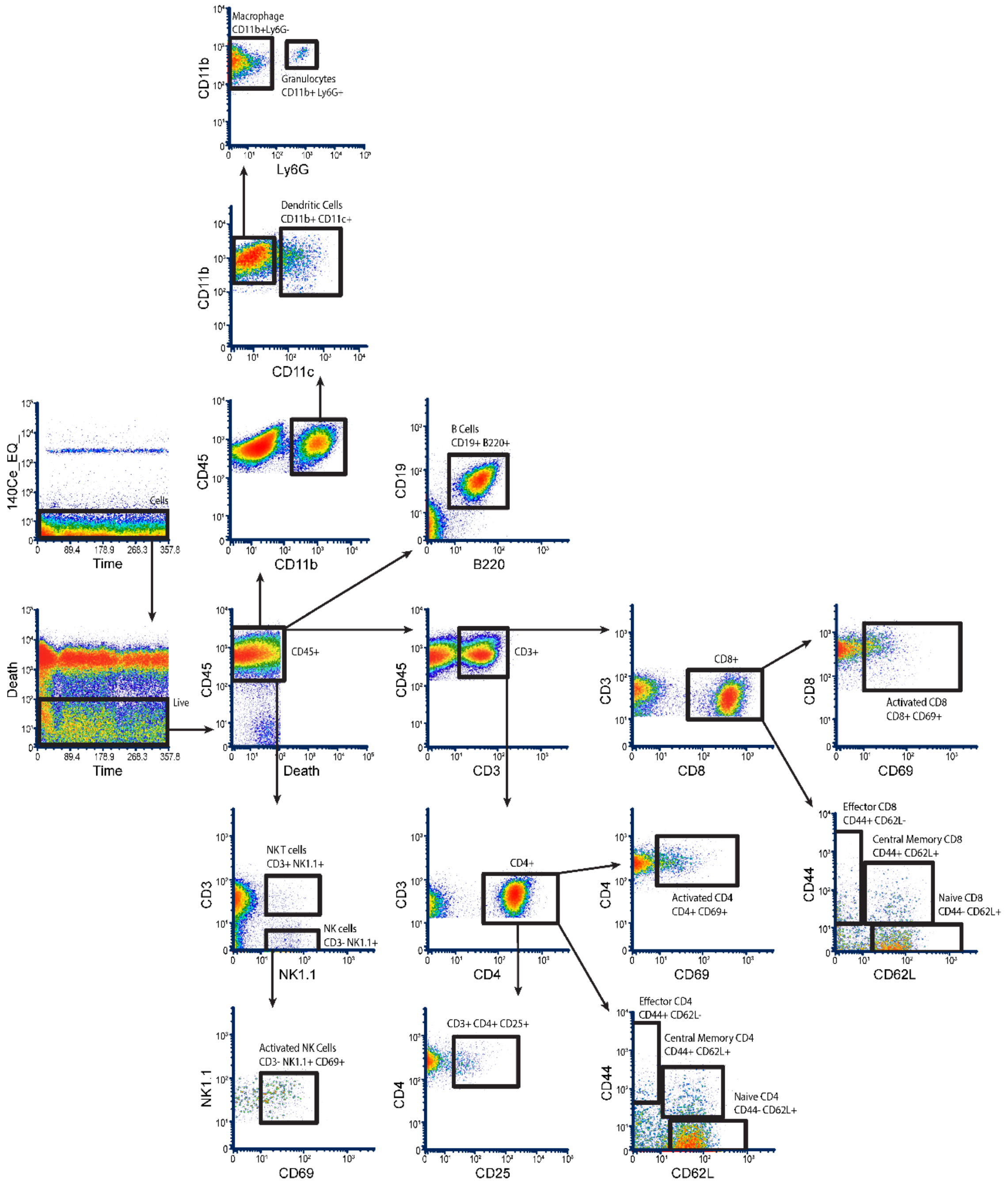


**Supplementary Figure S4:** Representative gating scheme for immune cell populations via cytometry by time of flight (CyTOF) of disassociated B16-F10 tumors in the flank of C57Bl/6 mice. A cascading gating strategy was established to first clean up the data by removing debris, aggregates, ion cloud fusions and dead cells and then differentiate leukocyte subpopulations based on marker expression. Populations of CD45$^+$ leukocytes further stratified into CD19$^+$B220$^+$ B cells, NK1.1$^+$TCRb$^+$ NKT cells, CD3$^+$TCRb$^+$ T cells (CD4$^+$ or CD8$^+$), NK1.1$^+$TCRb$^-$ NK cells, CD11b$^+$CD11c$^+$ dendritic cells, CD11b$^+$Ly6G$^+$ granulocytes and CD11b$^+$ myeloid cells. Activation marker CD69$^+$.

**Supplementary Video 1: Nanoparticle tracking analysis of extracellular vesicles - CHL-1 cells (siCtrl).**
Representative nanoparticle tracking analysis (NTA) of extracellular vesicles (EVs) from CHL-1 cells siCtrl via the NanoSight NS300. Individual nanoparticles were visualized under continuous flow and tracked to determine particle size distribution based on Brownian motion. Particle movement was recorded and analyzed using NTA software to calculate vesicle size and concentration.

**Supplementary Video 2: Nanoparticle tracking analysis of extracellular vesicles - CHL-1 cells (si*TPD52L2*).**
Representative nanoparticle tracking analysis (NTA) of extracellular vesicles (EVs) from CHL-1 cells si*TPD52L2* via the NanoSight NS300. Individual nanoparticles were visualized under continuous flow and tracked to determine particle size distribution based on Brownian motion. Particle movement was recorded and analyzed using NTA software to calculate vesicle size and concentration.

**Supplementary Video 3: C32 Melanoma VM formation (siCtrl).**
Time-lapse live-cell imaging of C32 melanoma cells seeded onto growth factor–reduced Matrigel demonstrating a dynamic vascular network formation over 24hrs post siCtrl transfection. Cells undergo alignment, elongation, and interconnection to form capillary-like structures characterized by branching points and enclosed loops. Imaging was performed using the CellVoyager CV1000 spinning disk confocal system. VM formation was defined as cellular incorporation and elongation forming connections between branch points, and was quantified by measuring total tube length, branching points, and loop number using ImageJ (v1.47).

**Supplementary Video 4: C32 Melanoma VM formation (si*TPD52L2*).**
Time-lapse live-cell imaging of C32 melanoma cells seeded onto growth factor–reduced Matrigel demonstrating a dynamic vascular network formation over 24hrs post si*TPD52L2* transfection. Cells undergo alignment, elongation, and interconnection to form capillary-like structures characterized by branching points and enclosed loops. Imaging was performed using the CellVoyager CV1000 spinning disk confocal system. VM formation was defined as cellular incorporation and elongation forming connections between branch points, and was quantified by measuring total tube length, branching points, and loop number using ImageJ (v1.47).

**Supplementary Video 5: CHL-1 Melanoma VM formation (siCtrl).**
Time-lapse live-cell imaging of CHL-1 melanoma cells seeded onto growth factor–reduced Matrigel demonstrating a dynamic vascular network formation over 24hrs post siCtrl transfection. Cells undergo alignment, elongation, and interconnection to form capillary-like structures characterized by branching points and enclosed loops. Imaging was performed using the CellVoyager CV1000 spinning disk confocal system. VM formation was defined as cellular incorporation and elongation forming connections between branch points, and was quantified by measuring total tube length, branching points, and loop number using ImageJ (v1.47).

**Supplementary Video 6: CHL-1 Melanoma VM formation (si*TPD52L2*).**
Time-lapse live-cell imaging of CHL-1 melanoma cells seeded onto growth factor–reduced Matrigel demonstrating a dynamic vascular network formation over 24hrs post si*TPD52L2* transfection. Cells undergo alignment, elongation, and interconnection to form capillary-like structures characterized by branching points and enclosed loops. Imaging was performed using the CellVoyager CV1000 spinning disk confocal system. VM formation was defined as cellular incorporation and elongation forming connections between branch points, and was quantified by measuring total tube length, branching points, and loop number using ImageJ (v1.47).

**Supplementary Video 7 – HT-144 Melanoma VM formation (siCtrl).**
Time-lapse live-cell imaging of HT-144 melanoma cells seeded onto growth factor–reduced Matrigel demonstrating a dynamic vascular network formation over 24hrs post siCtrl transfection. Cells undergo alignment, elongation, and interconnection to form capillary-like structures characterized by branching points and enclosed loops. Imaging was performed using the CellVoyager CV1000 spinning disk confocal system. VM formation was defined as cellular incorporation and elongation forming connections between branch points, and was quantified by measuring total tube length, branching points, and loop number using ImageJ (v1.47).

**Supplementary Video 8 – HT-144 Melanoma VM formation (si*TPD52L2*).**
Time-lapse live-cell imaging of HT-144 melanoma cells seeded onto growth factor–reduced Matrigel demonstrating a dynamic vascular network formation over 24hrs post si*TPD52L2* transfection. Cells undergo alignment, elongation, and interconnection to form capillary-like structures characterized by branching points and enclosed loops. Imaging was performed using the CellVoyager CV1000 spinning disk confocal system. VM formation was defined as cellular incorporation and elongation forming connections between branch points, and was quantified by measuring total tube length, branching points, and loop number using ImageJ (v1.47).

**Supplementary Video 9 – BMEC angiogenesis assay (siCtrl).**
Time-lapse live-cell imaging of BMEC endothelial cells seeded onto growth factor–reduced Matrigel demonstrating a dynamic vascular network formation over 24hrs post si*Ctrl* transfection. Cells undergo alignment, elongation, and interconnection to form capillary-like structures characterized by branching points and enclosed loops. Imaging was performed using the CellVoyager CV1000 spinning disk confocal system. Angiogenesis was defined as cellular incorporation and elongation forming connections between branch points, and was quantified by measuring total tube length, branching points, and loop number using ImageJ (v1.47).

**Supplementary Video 10 - BMEC angiogenesis assay (si*TPD52L2*).**
Time-lapse live-cell imaging of BMEC endothelial cells seeded onto growth factor–reduced Matrigel demonstrating a dynamic vascular network formation over 24hrs post si*TPD52L2* transfection. Cells undergo alignment, elongation, and interconnection to form capillary-like structures characterized by branching points and enclosed loops. Imaging was performed using the CellVoyager CV1000 spinning disk confocal system. Angiogenesis was defined as cellular incorporation and elongation forming connections between branch points, and was quantified by measuring total tube length, branching points, and loop number using ImageJ (v1.47).
